# Q-DOAS: A Proximity Quenching Assay for Real-Time Detection of Early Protein Aggregation Events

**DOI:** 10.64898/2026.06.16.732424

**Authors:** Korbin Michael Kleczko, Dan Gestaut, Sarah Dobbins, Juliana Abramovich, Cole Sitron, Xia Li, Rebecca Chan, Nan Wang, X. William Yang, F. Ulrich Hartl, Judith Frydman

## Abstract

Protein aggregation is central to numerous neurodegenerative diseases. The paucity of facile assays to measure aggregation and detect amyloid seeds hinders therapeutics development and clinical assessments. The standard ThT assay reports on amyloid formation but is blind to early oligomers and is prone to interference. We describe Q-DOAS, a simple and reproducible assay that quantifies aggregation in real time via proximity-quenching of a site-specifically conjugated dye (BODIPY-TMR). Using mutant Huntingtin-exon 1 (mHTT-Ex1) and α-Synuclein A53T, we show Q-DOAS detects pre-amyloid oligomers, yielding quantitative kinetic data compatible with mechanistic analysis. We further demonstrate its utility to dissect mutational effects, screen for protein and small-molecule aggregation inhibitors, and quantify amyloid seeding activity in cellular and mouse models of Huntington’s disease and in cerebrospinal fluid from Parkinson’s disease patients without amplification. Q-DOAS provides a sensitive, robust, and scalable tool for studying the earliest events in amyloid pathologies and for advancing therapeutic development.

## Introduction

Protein aggregation underlies a wide range of human diseases, including many neurodegenerative conditions. These diseases are characterized by formation of both oligomeric species and β-sheet-rich amyloid aggregates, which have been variously implicated in pathogenesis and disease spread^1,2^. The aggregation cascade proceeds from a monomer species through a succession of poorly characterized assemblies leading to amyloid aggregate formation^3–6^. The ability to assay and quantify the formation of both oligomers and amyloid or amyloid-like fibers (hereafter “amyloids”) in a time-resolved manner is crucial for understanding the mechanistic underpinnings and clinical progression of these diseases. The Thioflavin T (ThT) assay is the standard tool for quantifying aggregation and evaluating the effects of modulatory compounds, providing insight into the fundamental biology of amyloid-linked diseases and the efficacy of putative therapeutic approaches^7–9^. However, the utility of the ThT assay is limited because changes in fluorescence depend on the formation of β-sheet-rich amyloids, which appear to represent the terminal stage of aggregation^7–9^. Additionally, ThT can bind to a number of cellular protein assemblies as well as organic compounds that may limit its applicability in complex mixtures^10,11^.

It is increasingly recognized that soluble non-amyloid oligomers forming during the aggregation pathway can also have toxic functions in proteinopathies and amyloid diseases^12,13^. These small oligomeric species are not necessarily β-sheet-rich and are likely undetectable by the ThT assay unless they possess amyloid character^5,14–16^. This limitation has forced researchers to seek alternative methods for analyzing oligomer formation— ranging from Native PAGE to light-scattering, to FRET-based Analytical Size-Exclusion Chromatography — to elucidate the mechanisms underlying oligomer formation. Although useful, many of these methods lack the convenience and time-resolved resolution of the ThT assay, and most are relatively insensitive.

Using fluorescently labeled proteins could offer a more sensitive and time-resolved detection method for aggregation. A FRET-based assay was developed for polyQ-expanded HTT-Ex1 that is time resolved and sensitive^17^ but requires labeling of two proteins with distinct fluorophores and is susceptible to fluctuations in the immediate environment of the FRET probes. We sought to develop a more facile and versatile time-resolved fluorescence-based method to detect aggregation in real time by exploiting the intrinsic tendency of BODIPY-TMR and other fluorescent dyes to self-quench when in close proximity^18,19^. The dye was conjugated in a site-specific manner to the aggregation-prone protein of interest through maleimide labeling of a single cysteine at a location that does not interfere with aggregation, allowing dynamic monitoring of intermolecular interactions by proximity quenching. This approach enabled time-resolved detection of oligomer and aggregate formation in a plate reader-based assay. Because aggregation is a seeded process, the assay can also detect and quantify the presence of amyloid seeds in biological samples. In addition, the effect of proteins and reagents that would otherwise interfere with ThT fluorescence can be tested without confounding the readout, while retaining the plate reader format.

We here developed the Q-DOAS (Quantitative Detection of Oligomer and Amyloid Seeds) assay and demonstrated its application to detect oligomerization and aggregation of two amyloidogenic proteins commonly studied using ThT: mutant Huntingtin exon 1 (mHTT-Ex1) carrying a pathogenic expansion in its polyQ tract and Parkinson’s Disease linked mutation in α-Synuclein A53T (αSyn A53T). We benchmark the approach against traditional biochemical detection methods, including native gel, filter retardation trap and ThT, and demonstrate its utility to measure the effect of mutations, aggregation inhibiting protein and small molecule ligands. Finally, we demonstrate Q-DOAS’ ability to detect amyloid seeds in biological samples, ranging from CSF to mouse brain lysates. The facile, quantitative and time-resolved nature of the Q-DOAS assay should make its adaptation possible to examine a broad range of aggregation and amyloid processes.

## Results

### Q-DOAS assay principle and kinetic readout

We reasoned that fluorescence proximity quenching can be exploited to follow aggregation kinetics in real time for a range of amyloid disease linked proteins (Figure 1A). Fluorescence self-quenching by proximity is a phenomenon where the signal of a fluorophore is dramatically reduced due to close physical contact with another dye or quenching molecule. Dyes like TMR (Tetramethylrhodamine) and BODIPY (borondipyrromethene) are frequently combined to build this “turn-off/turn-on” mechanism. When BODIPY-TMR is in close contact, energy transfer is nearly 100% efficient, effectively turning off the emission^20^. When the protein of interest is labeled with BODIPY-TMR, it will fluoresce in the monomeric state, but fluorescence will be quenched during oligomerization and aggregation. This approach enables the time-resolved quantification of aggregation kinetics. As kinetic data of protein aggregation can be fitted to infer molecular mechanism for aggregation reactions^3^, we reasoned the Q-DOAS assay could open the way for obtaining mechanistic insights into a wide range of protein pathologies, as well as to examine approaches to inhibit their aggregation from the earliest events in the aggregation pathway.

**Figure 1.**
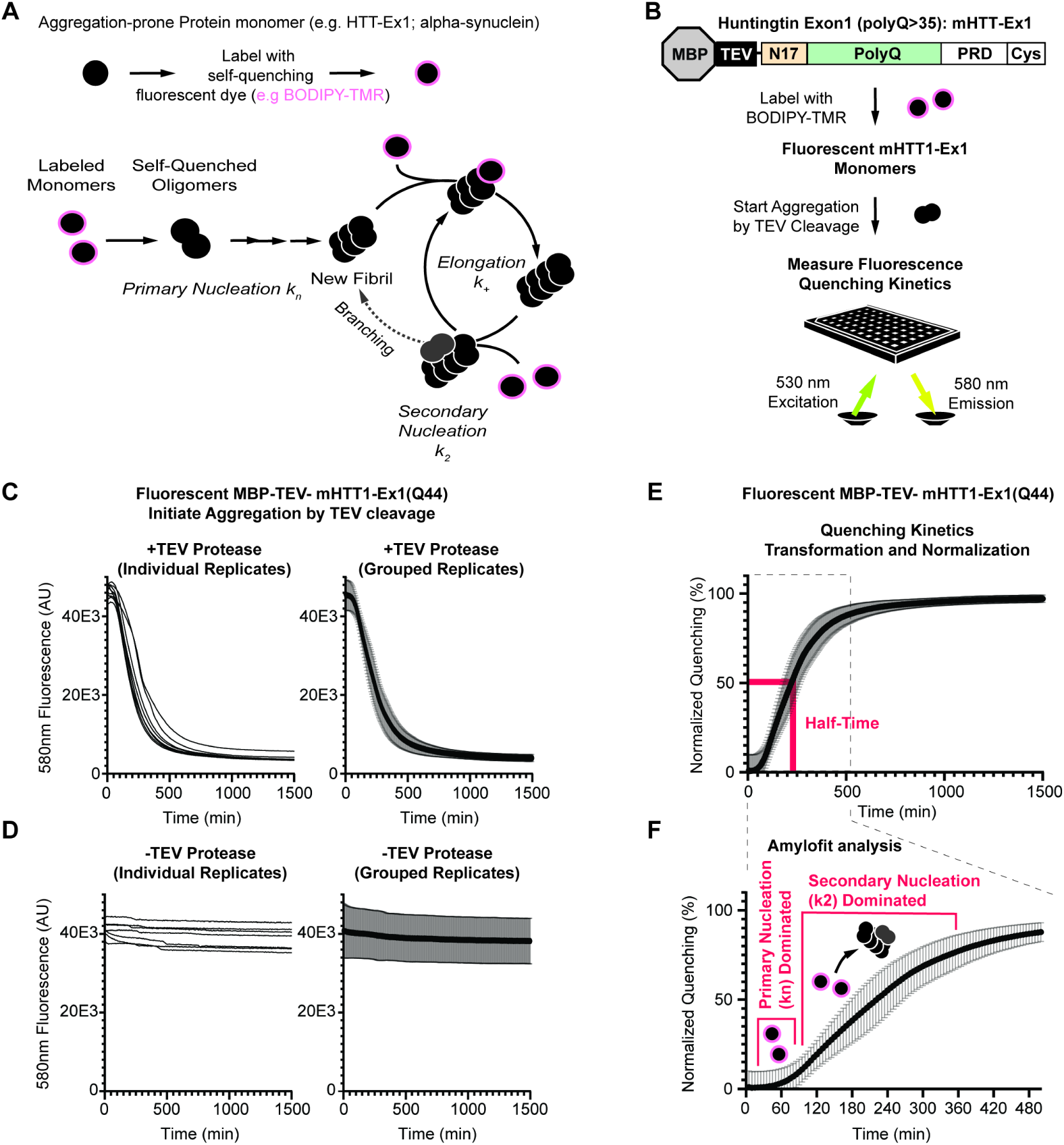
Q-DOAS assay principle and kinetic readout: Development for polyQ-expanded HTT-Ex1. Assay shown represents one assay with n=9 replicates. (A) Schematic representation of MBP-mHTT-Ex1 undergoing aggregation, especially as it pertains to plate-reader based assays^3,4^. A C-terminal cysteine is used for specific labeling of a fluorescent and self-quenching moiety. Primary nucleation (*k_n_*) is a slower process seeding aggregation, whereas elongation (k+) is assumed to be constant. Secondary nucleation (k2) events increase the overall amount of aggregation, increasing seeding and elongation events as a whole, which can be observed in Q-DOAS. (B) General schema from protein to assay using a mHTT-Ex1-BDP example. Unlabeled mHTT-Ex1 is labeled using BDP-TMR, samples are added to a 384-well plate, and aggregation is induced through release from the soluble MBP moiety and fluorescence is measured at regular intervals. (C) Q-DOAS using 1.5 µM MBP-mHTT-Ex1(Q44)-BDP in the presence of 1 µM uTEV3. The left graph shows individual replicates and the right graphs shows the average and standard deviation. The assay was allowed to proceed until all replicates showed no observable changes in quenching. (D) Q-DOAS using 1.5 µM MBP-mHTT-Ex1(Q44)-BDP without uTEV3. The left graph shows individual replicates and the right graphs shows the average and standard deviation. Samples exhibit no significant changes in fluorescence quenching without releasing the sample from MBP. Similarly, the labeled protein must be aggregation-prone to report aggregation. (E) Q-DOAS results from (C) have been transformed about the *x*-axis and normalized to adopt a more familiar conformation for ease of observation. Normalization allows for direct comparison between samples which exhibit slight differences in overall fluorescence intensity. (F) Normalized Q-DOAS results from (E) are shown here with a focus on the first 8 hours of aggregation. The initial lag phase is thought to be primary nucleation-dominated before secondary nucleation accelerates the rate of quenching, as seen in the graph results.

### Developing Q-DOAS for polyQ-expanded HTT-Ex1

Wild-type huntingtin (HTT) is a 348 kDa protein implicated in neuronal vesicle trafficking^21–23^. Expansion of the polyglutamine (polyQ) tract in its exon 1 beyond 35 repeats renders it aggregation prone, giving rise to neurotoxic species that lead to Huntington’s Disease (HD)^1,13^. In HD, aggregates contain predominantly short forms of HTT, predominantly comprising HTT Exon 1 (HTT-Ex1), which by itself suffices sufficient to induce disease phenotypes in patients and to recapitulate amyloid-forming aggregation *in vitro* and *in vivo*^24–27^. HD studies typically employ mHTT-Ex1 to understand its *in vitro* modulation, or to examine aggregation and toxicity in animal or cellular models^13,17,28,29^. While the polyQ tract of HTT-Ex1 length drives aggregation propensity, its two flanking sequences, the N-terminal N17 amino acids, called N17, and its C-terminal proline-rich domain (PRD) modulate its aggregation pathway^5,30–32^ (see below Fig. 4). Approaches to study HTT-Ex1 aggregation *in vitro* primarily rely on amyloid detection approaches which do not detect early oligomerization events.

To implement the Q-DOAS assay for real-time observation of oligomer and aggregate formation we employed an mHTT-Exx1(Q44) maintained soluble via fusion to Maltose Binding Protein (MBP) that is cleaved by Tobacco Etch Virus Protease (TEV) to initiate aggregation. BODIPY-TMR was conjugated to MBP-mHTT-Ex1(Q44) via maleimide chemistry to a C-terminal Cysteine to generate MBP-mHTT-Ex1-BDP (Figure 1B). This labeling site was selected because the PRD is not incorporated into the amyloid fibril structure but instead projects outward from the polyQ-rich cross-β core^31,33–35^; attaching BODIPY-TMR to a flexible, solvent-exposed region facilitates dye–dye proximity upon aggregation while reducing the likelihood of interference with aggregation-promoting interactions^10^. While the chosen labeling site was previously shown to be compatible with HTT-Ex1 aggregation^36–38^, we also tested various ratios of labeled MBP-mHTT-Ex1-BDP to unlabeled MBP-mHTT-Ex1 and chose to work with a sub-stoichiometric ratio of labeled MBP-mHTT-Ex1-BDP (Extended Data 1A,C-E). The reaction is initiated by addition of TEV protease in the assay plate to cleave MBP from mHTT-Ex1-BDP (Figure 1B). Samples are incubated at 30°C and measured every 5 minutes to capture changes in fluorescence on a plate reader (Figure 1B), reported in arbitrary units (AU). When TEV protease was added, the time-resolved measurements of fluorescence intensity at 580 nm decayed with characteristic and highly reproducible kinetics (Figure 1C). Importantly, omitting TEV, which abrogates aggregation, also prevented any changes in fluorescence (Figure 1D), indicating that proximity quenching results from the HTT-Ex1 aggregation process.

In order to facilitate fitting the fluorescence decay kinetic analyses using existing platforms, such as Amylofit, we normalized and transformed our measurements using the formula:

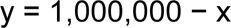

where x is the measured fluorescence intensity in AU. The arbitrary constant 1,000,000 was chosen to be safely above the maximum dynamic range of the plate reader, ensuring that the inversion yields positive values across the full signal window. Transformed values are then normalized by setting the minimum transformed value to 0 and the maximum to 1. This inversion and normalization convert the data into a format that is readily amenable to quantitative analysis platforms without altering the underlying result (Figure 1E). The resulting curves can be analyzed to determine half times, lag times, and nucleation constants using established fitting procedures such as Amylofit (Figure 1F). This curve can be interpreted analogously to other aggregation-based assays, such as ThT, to track changes in mHTT-Ex1-BDP quenching over time^2–4,39^. We observe that the HTT-Ex1(Q44) aggregation kinetics contain a lag phase with little changes in quenching, whose duration is determined by primary nucleation kinetics (Figure 1F). Upon production of sufficient nuclei, quenching increases as available monomers are depleted into oligomers and aggregates. The rate of the slope is linked to secondary nucleation rates. Finally, the reaction reaches a plateau with no further changes in signal that can be used to calculate the half time of the aggregation reaction. The assay cannot define the precise structural state of quenched mHTT-Ex1-BDP; rather, it serves as a proxy for the rate at which oligomerization and aggregation events occur and the relative aggregation propensity under these experimental conditions.

### Biochemical mapping of the quenching signal for mHTT-Ex1 Q-DOAS

To better understand the underlying biochemical processes leading to the Q-DOAS readout, we biochemically characterized aggregate species formed at selected time points. To this end, we carried out two identical aggregation reactions using 1.5 µM mHTT-Ex1-BDP in Reaction Buffer A at 30°C. One was used for the Q-DOAS assay and the other provided samples for biochemical analyses (Figure 2A). Time points were selected to capture inflection points of interest along the Q-DOAS quenching curve. Samples were snap-frozen in liquid nitrogen (LN₂) to arrest further aggregation until analysis (Figure 2B). HTT-Ex1 species were detected using an mAB5492 (anti-PRD) immunoblot.

**Figure 2.**
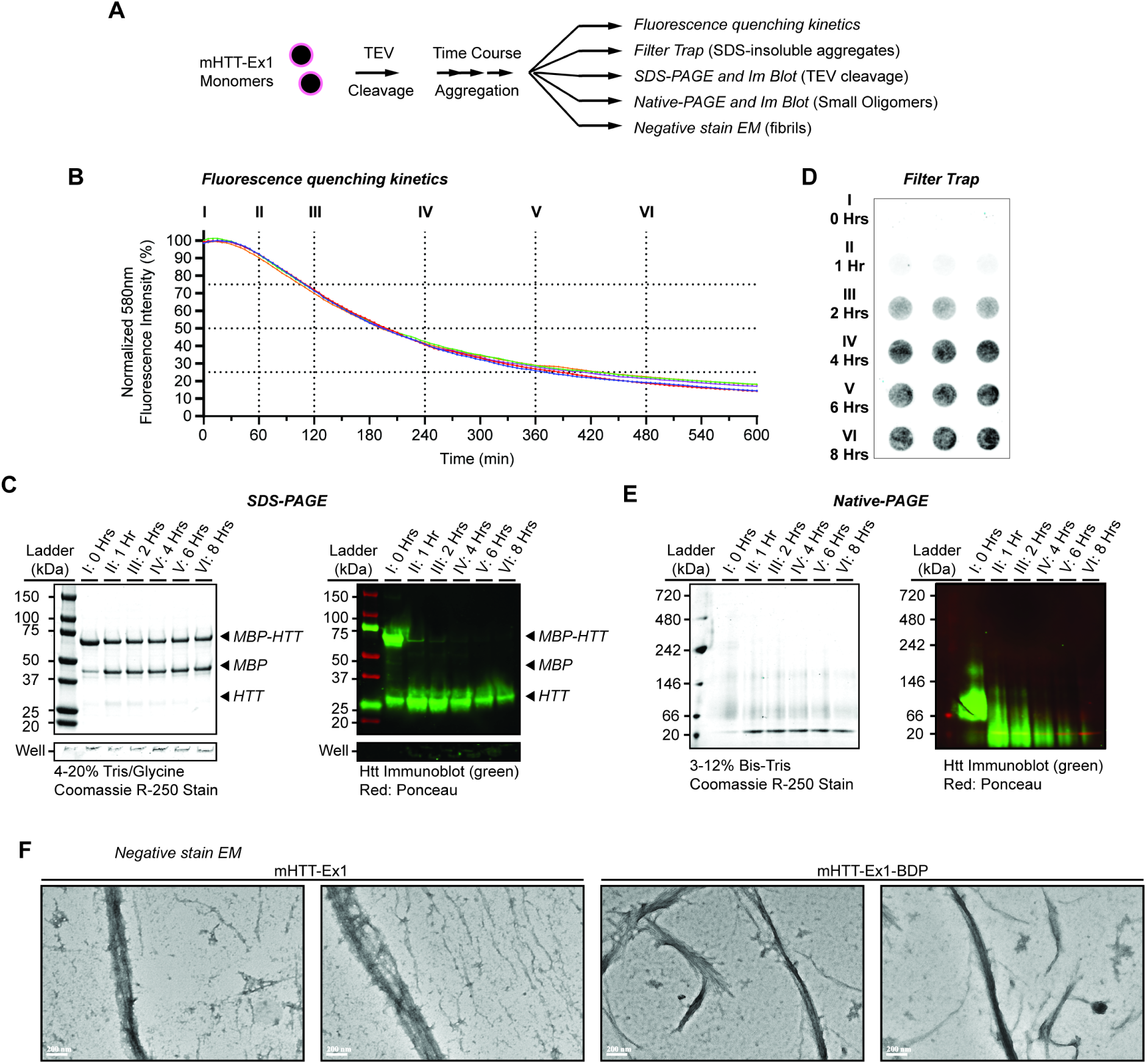
Biochemical mapping of the quenching signal for mHTT-Ex1 Q-DOAS. (A) A singular mHTT-Ex1-BDP master mix was prepared to perform each experiment in parallel. Samples were permitted to aggregate before analysis using different methods. (B) Representative Q-DOAS assay with individual replicates shown. Individual replicates under controlled experimental conditions show little variance between replicates. A C-terminal cysteine is used for specific labeling of a fluorescent and self-quenching moiety. Assay displays n=9 technical replicates. (C) SDS-PAGE gel tracking the depletion of SDS-soluble species. Left gel shows all protein using coomassie R-250; right membrane is the corresponding Western Blot using anti-HTT antibodies. Assay displays n=1 replicate. (D) FT assay in triplicate showing amyloid formation over the time course. Assay displays n=3 technical replicates. (E) BN-PAGE gel tracking the depletion of oligomeric species. Left gel shows all protein using coomassie R-250; right membrane is the corresponding Western Blot using anti-HTT antibodies. Assay displays n=1 replicate. (F) Negative Stain Electron Micrograph images of mHTT-Ex1 and mHTT-Ex1-BDP show high visual similarity of fibril structure and bundling behavior.

SDS-PAGE indicated the TEV cleavage reaction begins to release mHTT-Ex1 from MBP at the earliest timepoint analysed (approx. 5 min) and is fully complete at 1 hour (Figure 2C). Monomeric mHTT-Ex1-BDP was depleted over time, with SDS-insoluble aggregates visible in the well of the SDS-PAGE gel at later time points (Figure 2C). We also used the Filter Trap (FT) assay to track formation of SDS-insoluble amyloids^40^. While fluorescence quenching indicated formation of oligomeric species at 1 hour in the Q-DOAS assay, FT started detecting amyloids only by two hours of assay initiation (Figure 2D), and reached a plateau at 4 hours (Figure 2D). This suggests SDS-insoluble amyloids alone do not fully account for the observed quenching signal. Blue Native-PAGE revealed the presence of smaller oligomeric mHTT-Ex1 species, peaking at 1-2 hours before declining (Figure 2E), consistent with a transition from oligomers to amyloids during the time course. These observations agree with current nucleation–elongation models of mHTT-Ex1 aggregation. While Q-DOAS cannot define the structural state of mHTT-Ex1-BDP at any given time point, persistent dye–dye interactions provide a proxy for both the rate of oligomerization and aggregation and the relative aggregation propensity across experimental conditions.

### Generalization to αSyn and comparison to ThT

Real-time tracking of αSynuclein (αSyn) oligomerization and aggregation presents distinct challenges relative to mHTT-Ex1. The slower kinetics of unseeded αSyn aggregation and lower aggregation propensity of αSyn compared to HTT-Ex1, combined with the need for regular agitation, necessitates longer measurement intervals to capture the full progression from lag phase to plateau, reducing the overall temporal resolution of these measurements. ThT is the most established method for studying αSyn aggregation^41–43^ and allows direct detection of amyloids. However, ThT cannot detect non-amyloid species and its fluorescence is sensitive to changes in pH and interference by biological macromolecules^42^. We sought to develop a Q-DOAS assay for αSyn and compared it to the ThT standard. To this end, we compared ThT and Q-DOAS to monitor aggregation of αSyn A53T, one of the primary genetic drivers of familial Parkinson’s disease (PD). This autosomal dominant missense mutation dramatically accelerates protein misfolding and aggregation, driving early-onset parkinsonism^44,45^. BODIPY-TMR labeling was achieved through maleimide chemistry by introducing a single cysteine at position A140; previous findings showed this modification minimally alters αSyn aggregation kinetics when substoichiometrically labeled^46^ (Figure 3Ai, Extended Data 1B). Since unseeded αSyn aggregation is extremely slow^47^, we also tested the ability of both assays to assess seeded aggregation. αSyn seeds generated from unlabeled monomers were prepared using established protocols^48^. Seeds were snap-frozen in LN₂ and thawed at room temperature before addition to the assay (Figure 3Aii). Seeds were snap-frozen in LN₂ and thawed at room temperature before addition to the assay.

**Figure 3.**
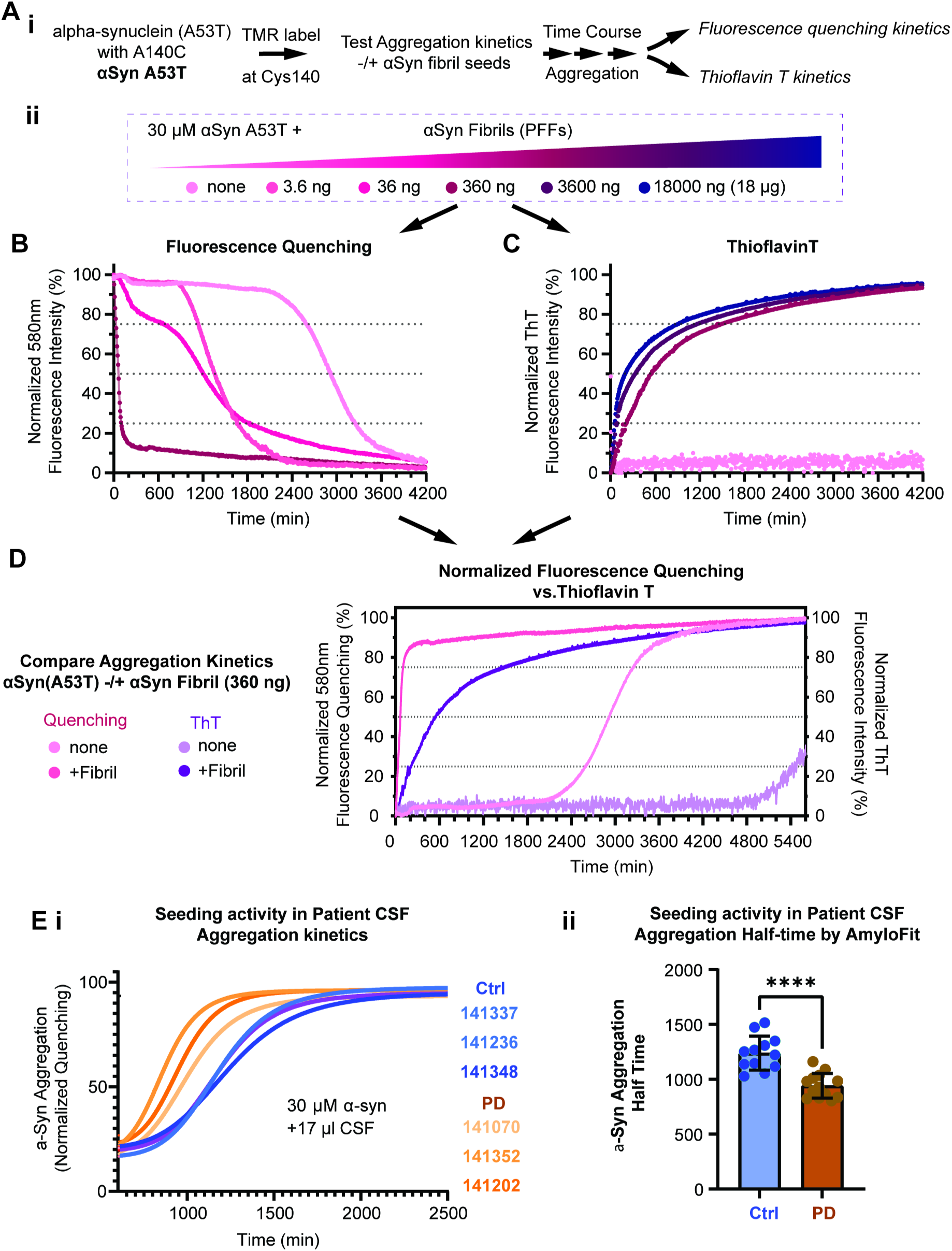
Developing Q-DOAS for αSynuclein and comparison to ThT. (A) The A140C mutation on the ɑSyn A53T monomers is located C-terminal to the xKTEGVxxx repeats to avoid interfering with the formation of the amyloid fibril. (Ai) ɑSyn A53T was allowed to aggregate in the presence of increasing concentrations of fibrils and analyzed either using Q-DOAS or ThT. Monomers and fibrils are loaded onto the plate at RT and placed in the plate reader with shaking according to the protocol to induce aggregation for both Q-DOAS and ThT. No beads or additional additives are used to induce aggregation. (Aii) ɑSyn A53T PFF concentration began at 3.6 ng, increasing to 18 µg at the highest. (B-C) Both Q-DOAS (B) and ThT (C) curves exhibit quenching of seeded aggregates at around 4200 minutes. However, only Q-DOAS sees complete quenching of ɑSyn(A53T)-BDP by the 4200 minute mark, suggesting the formation of a non-amyloid aggregate which is not visible through ThT. Q-DOAS curves display the average of n=3 replicates. ThT curves display the average of n=4 replicates. (D) Overlaid Q-DOAS and ThT assay curves using equivalent fibril concentrations show distinct behavior. Most notably, we observe a 600 minute delay between complete Q-DOAS quenching of unseeded ɑSyn(A53T)-BDP and the start of unseeded ɑSyn(A53T) ThT fluorescence. (E) Quantifying seeding activity in patient CSF. Proof of concept that the seeding activity in patient CSF can be differentiated using ɑSyn(A53T)-BDP. (Ei) Grouped datapoints show CSF from PD+ patients exhibit faster aggregation curves compared to PD-controls. Each curve represents the average of n=5 replicates. (Eii) Half times of Control vs. PD+ patients show a significant decrease in PD+ half times (Mean Diff.: 297.3, 95% CI of Diff.: 123.5 to 471.2)(one-way ANOVA: F = 8.905, P <0.0001; CTRL vs PD+ P <0.0001; PD-n=12, PD+ n=13).

Q-DOAS (Figure 3B) and ThT (Figure 3C) assays were carried out in parallel using the same monomer and seed concentrations and time-course. It is immediately apparent that the Q-DOAS assay detects oligomers and aggregates earlier in the time course and responds to seeding with higher sensitivity than ThT. For instance, in the absence of seeds, there was no detectable αSyn aggregation signal in the ThT assay until at least 80 hrs, while the Q-DOAS assay showed quenching with a lag phase of 33 hrs. Q-DOAS also responded more sensitively and dynamically to seed concentrations (Figure 3B, 3C), suggesting either this initial lag phase, or the half-time of fluorescence decay may serve as metrics for estimating the quantity of seeding-competent material in a sample. Direct comparison of the normalized kinetics of αSyn aggregation by Q-DOAS or ThT without or with 360 ng PFF seeds (Figure 3D) demonstrates Q-DOAS detects the formation of oligomers and aggregates significantly earlier than the ThT assay (Figure 3D). The earlier quenching signal is consistent with formation of oligomers and aggregate species invisible to ThT. Additionally, the Q-DOAS assay plateaued earlier than ThT, suggesting αSyn forms structurally immature aggregates before adopting the cross-β configuration required for ThT binding.

### Quantifying seeding activity in patient CSF (proof-of-concept)

There are currently many established and emerging methods for amyloid disease diagnosis^9,49^, but most rely on detection of ThT-reactive amyloid seeds. Q-DOAS offers an orthogonal approach in which pre-amyloid seeding activity can be tracked. We next examined the ability of Q-DOAS to detect αSyn seeds in the CSF of Parkinson’s Disease patients (Figure 3E). αSyn concentration in CSF is extremely low, ranging from 0.2-2 ng/ml. Given the insensitivity of ThT, the presence of seeds in PD CSF is detected via Seed Amplification Assay (SAA) to detect the minute fraction of seeds. The SAA is traditionally binary (positive/negative), and of limited use to predict cognitive decline stage or measure the effectiveness of disease-modifying therapies^50^. We used CSF from PD and control (Ctrl) patients previously characterized using the SAA assay^49^. The Q-DOAS study was carried out in quadruplicate with 17 µL of complete patient CSF in a double-blind experiment (i.e. the PD-status of the patient was blinded until after the experiment). There were clear differences in the aggregation kinetics between the CSF of PD patients and non-PD controls (Figure 3Ei). This was confirmed when we calculated the half-times of aggregation using the Amylofit data analysis pipeline to fit the kinetic data (Figure 3Eii). These experiments indicate that Q-DOAS has enough sensitivity to detect the very low αSyn concentrations in PD serum (∼3.5-35 pg) even without any amplification of capture, highlighting its potential for therapeutic quantification of αSyn seeds in patient samples.

### Dissecting HTT flanking-region effects on seeded kinetics using Q-DOAS

N17 and the PRD have been shown to affect HTT-Ex1 aggregation propensity, level of oligomer formation and aggregate morphology (Fig. 4A)^6,30,32,51^. The quantitative and time-resolved nature of Q-DOAS offered the opportunity to examine the effect of N17 and PRD flanking regions both on the ability to seed monomers or the ability of mutant monomers to become incorporated into fibrils (Figure 4). To this end, we generated BODIPY-TMR labeled variants of WT, ΔN17 and ΔPRD mHTT-Ex1(Q44), bearing a TEV-cleavable MBP tag. Upon TEV cleavage, the unseeded Q-DOAS aggregation half-time results behaved as expected from previous studies^52–54^ with ΔN17 aggregating much more slowly than WT and ΔPRD aggregating much faster.

**Figure 4.**
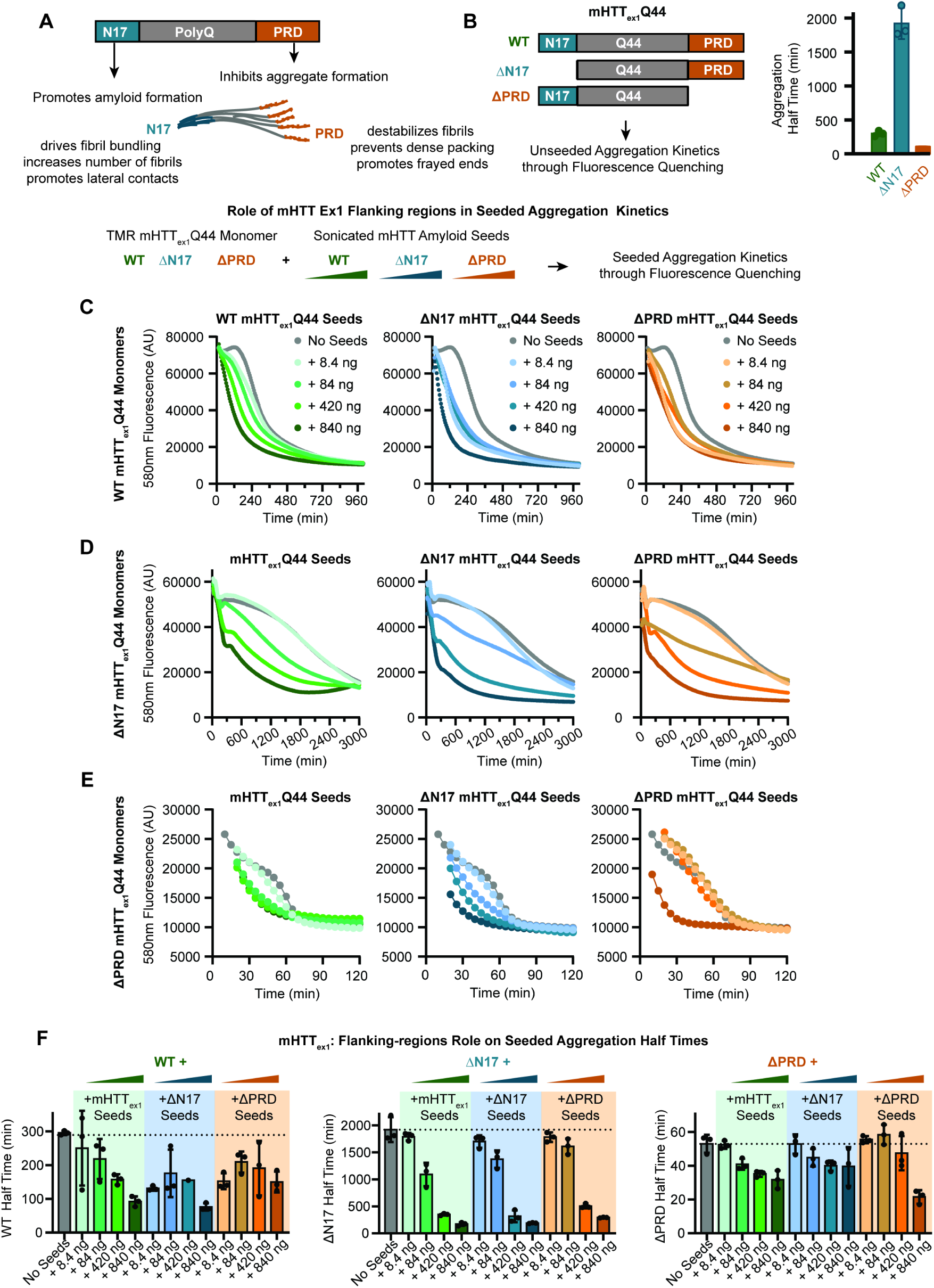
Dissecting HTT flanking-region effects on seeded kinetics using Q-DOAS. (A) Representation of the mHTT-Ex1 monomer variants used for this study. BDP-TMR was C-terminally conjugated to a cysteine (not shown) on each construct. (B) Abbreviated workflow for the formation and addition of amyloid fibrils to assay conditions. Sonicated seeds are added to monomeric mHTT-Ex1(Q44) variants in increasing concentrations from 0 ng to 840 ng per well. Bar graph exhibits expected unseeded aggregation behavior from mHTT-Ex1(Q44)-BDP, ΔN17(Q44)-BDP, and ΔPRD(Q44)-BDP (n=3 replicates per condition). (C-E) Cross seeding Q-DOAS experiments using 1.5 µM of free BDP-labeled monomers and variable amounts of amyloid fibrils. (C) Curves showing averaged Q-DOAS replicates in the presence of mHTT-Ex1(Q44)-BDP with different seeds (n=3 replicates per condition). (D) Curves showing averaged Q-DOAS replicates in the presence of ΔN17(Q44)-BDP with different seeds (n=3 replicates per condition). (E) Curves showing averaged Q-DOAS replicates in the presence of ΔPRD(Q44)-BDP with different seeds (n=3 replicates per condition). (F) Half times for each curve seen in C-E. Derived half times were plotted for direct comparison between each condition. Half times using different BDP-labeled monomers are listed on the *y*-axis (n=3 replicates per condition).

To examine the effect of N17 and PRD on seeded aggregation, we next generated aggregates for each unlabeled HTT-Ex1 variant. Aggregates were sedimented and then sonicated to generate seeds exactly as described in Shen et al. (2016)^30^. Increasing concentrations of each type of HTT-Ex1 seeds (i.e. WT, ΔN17 and ΔPRD) were used to seed the different monomer variants (Figure 4C-E). Regardless of the monomer variant and the type of seed, we observed a seed-concentration dependent reduction in the initial lag phase of aggregation and the half-time of fluorescence quenching (Figure 4F). Strikingly, the overall rate and extent of aggregation appeared determined primarily by the monomeric HTT-Ex1 variant used as substrate in the assay (Figure 4C–F). WT HTT-Ex1(Q44) aggregation was enhanced approximately 3-fold by WT, ΔN17 and ΔPRD seeds, from approximately 300 min half-time to ∼50-100 min at the highest seeds concentrations (Figure 4C, 4F). Seeding had the most dramatic effect on ΔN17 HTT-Ex1(Q44) (Figure 4D, 4F). Its unseeded half-time was approximately 1,900 min (approximately 31.6 hours), which was reduced to 171 minutes at the highest concentration. In contrast, the fast ΔPRD aggregation was least sensitive to seed addition, with a half time of 50 min for the unseeded reaction reduced to approx 20-30 minutes with ΔPRD and WT seeds and virtually unaffected by ΔN17 seeds. These experiments indicate that the flanking regions play a complex role in formation of aggregation competent monomer conformations as well as in primary nucleation. In particular, the very rapid aggregation of ΔPRD and its relative insensitivity to seed addition suggests that primary nucleation is not rate limiting for this variant. On the other hand, the very high sensitivity of ΔN17 to seeding suggests it is unable to form amyloid states that are seed competent. This feature makes it an attractive substrate to detect amyloid-seeding competent species of pathogenic HTT in biological samples.

### mHTT-Ex1 Q-DOAS Assay sensitivity to pH and salt concentration

We next examined the effects of buffer, salt concentration and pH on mHTT-Ex1 Q-DOAS aggregation kinetics. The charged residues in N17 and the PRD flanking domains of mHTT-Ex1 can adopt different conformations and interactions depending on pH, salt and solvent conditions^30,55^. pH-dependent conformational changes can influence the seeding and aggregation potential of amyloid-linked proteins. Low pH has been shown to reduce aggregation, and promote helical conformations in mHTT-Ex1, with progressive loss of helical character as pH increases toward alkaline conditions^55–57^. Elevated salt concentrations may also confound the evaluation of reagents that rely on electrostatic interactions with mHTT-Ex1. We asked whether distinct conformations and interdomain associations are detectable by Q-DOAS.

We used Q-DOAS to examine the effect of salt concentration and pH on the unseeded aggregation kinetics of WT, ΔN17 and ΔPRD HTT-Ex1(Q44). Salt concentration was varied from 100 mM to 500 mM NaCl. WT mHTT-Ex1 showed modest changes in lag phase and slope with increasing salt (Extended Data 2A), consistent with a role for ionic interactions in the kinetics of full-length construct aggregation. In contrast, ΔN17 aggregation showed little salt-sensitivity, suggesting that its aggregation kinetics are not strongly dependent on ionic interactions (Extended Data 2A). ΔPRD aggregation was also salt insensitive, though its rapid aggregation rate makes precise assessment of variability difficult.

Changes in pH most profoundly affected aggregation behavior, in some cases qualitatively altering the shape of the Q-DOAS curve. WT mHTT-Ex1 aggregation at pH 5.0 was markedly delayed, suggesting reduced conformational favorability at acidic pH, consistent with prior studies^55–57^ (xtended Data 2B). This inhibitory effect diminished progressively as pH was raised from 6.8 to 8.0. WT mHTT-Ex1 exhibited a more pronounced and multiphasic response, displaying a shortened initial lag phase and a gradual reduction in slope as pH increased (Extended Data 2B). In contrast, ΔN17-BDP and ΔPRD-BDP curves increased consistently across this pH range. This multiphasic behavior —absent in either flanking region truncation— is consistent with the hypothesis that the N17 and PRD flanking domains compete to modulate polyQ aggregation propensity, an effect that is unmasked only when both domains are present^58–61^.

### Screening aggregation modifiers of HTT using Q-DOAS

We next applied Q-DOAS to evaluate putative mHTT aggregation modifiers (Figure 5A). mHTT-Ex1 association to proteins and membranes can modulate its aggregation pathway — e.g. through membrane-induced nucleation or chap^52,62,63^ mediated suppression^64–66^. These events have been difficult to characterize using ThT. For instance, many chaperones bind to ThT, and their effect on amyloid formation must be monitored by less sensitive methods such as the FT assay The recent emergence of protein-based therapeutics targeting mHTT-Ex1 pathology has created a need for assay methods that are scalable and reliably quantifiable.

**Figure 5.**
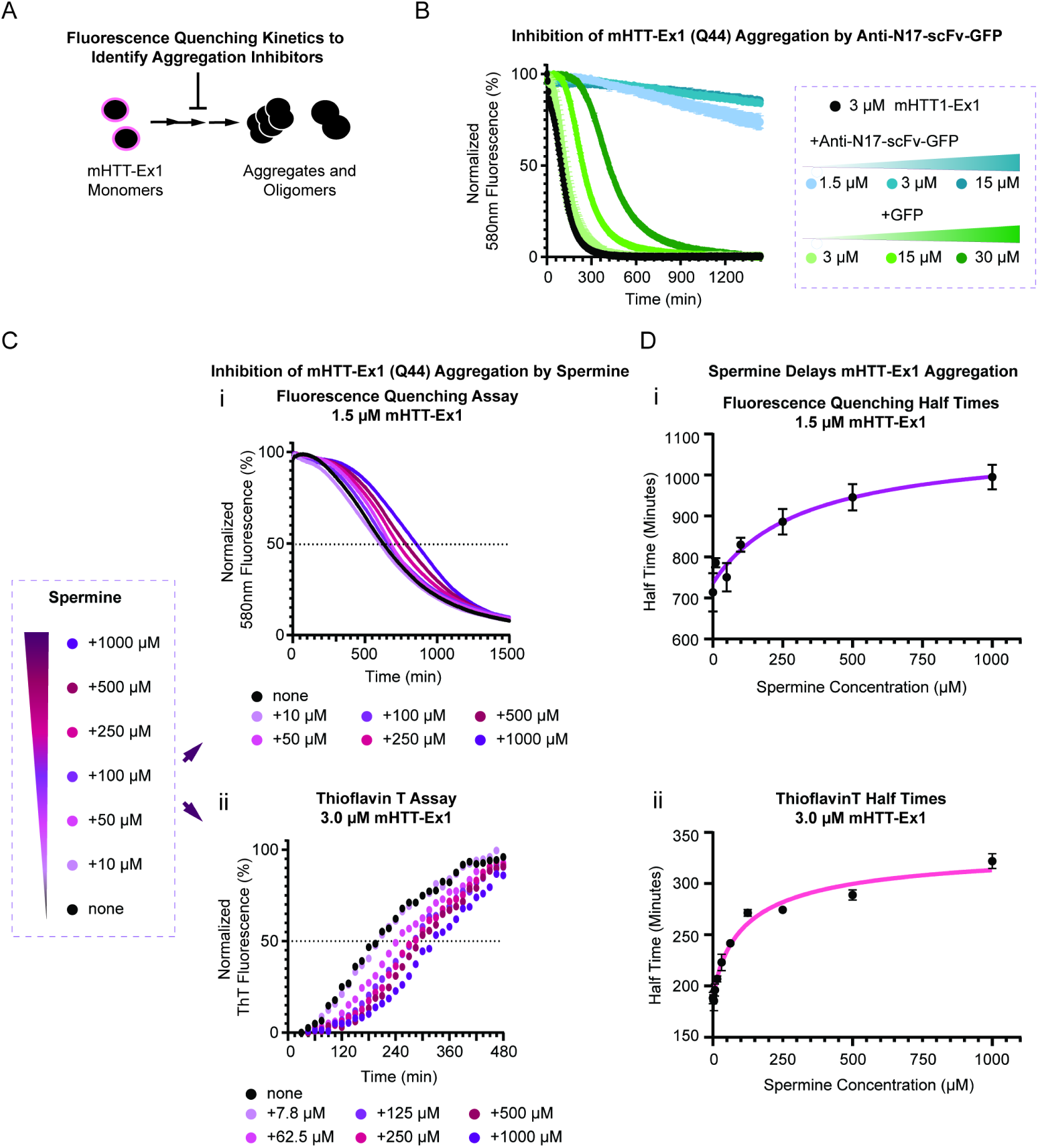
Screening aggregation modifiers of HTT using Q-DOAS. (A) Schematic representation of Q-DOAS in the presence of aggregation inhibitors. Aggregation suppression is measured via the persistence of fluorescence over an inhibitor-free control to determine efficacy. (B) αN17-GFP and GFP alone were tested for dose-response behavior using protein-based aggregation modifiers in the presence of 3 µM mHTT-Ex1(Q44)-BDP. Aggregation half times were recorded and plotted using a bar graph (Extended Data 3A). Notably, αN17-GFP was not observed to induce aggregation in the testing period (No Aggregation, “NA”), and has not been recorded here. Extended testing periods may show aggregation at later time points not collected in this experiment. Data shown represents the average of n=2 replicates. (C) Varying contractions of free spermine in solution were added to either (i) a Q-DOAS experiment using 1.5 µM mHTT-Ex1(Q44)-BDP or (ii) a ThT experiment using 3.0 µM mHTT-Ex1(Q51) and varying contractions of free spermine in solution. Q-DOAS curves represent the average from n=3 replicates. ThT curves represent the average from n=2 replicates. (D) Dose-response curve showing half-times from 5C derived from either the Q-DOAS Assay (i) or the ThT Assay (ii).

We first tested aggregation suppression using VL12.3, a single-chain variable fragment (scFv) intrabody that targets the N17 domain^67^. VL12.3 has previously been shown to suppress amyloid formation and toxicity^67^. We fused VL12.3 to GFP (herein anti-N17-scFv -GFP) 48 to improve its solubility; unconjugated GFP was included as a control for any independent effects of GFP on HTT-Ex1 aggregation (Figure 5B). Time resolved fluorescence quenching over a 25-hour period, showed anti-N17-scFv -GFP effectively suppressed HTT-Ex1 aggregation, even at a sub-stoichiometric molar ratio (Figure 5B), consistent with previous reports. At molar ratio or when in excess, the inhibition of aggregation was nearly complete. Interestingly, while the GFP control had virtually no effect on HTT-Ex1 aggregation at concentrations equimolar to HTT-Ex1, it has an inhibitory effect on HTT-Ex1 aggregation at high concentrations (Figure 5B). The increase in aggregation half-time was GFP concentration dependent (Extended Data 3A), indicating it is mediated by a low affinity GFP interaction. While GFP is commonly used as a fluorescent reporter to visualize and quantify mHTT-GFP aggregation^12,65^, there are reports that fusion to GFP changes the nature of HTT-Ex1 aggregates^68,69^. Although these findings do not invalidate prior results using mHTT-GFP fusions, they suggest that mHTT-Ex1 — and potentially other amyloidogenic proteins — can interact with GFP in ways that may subtly influence outcomes in certain experimental contexts.

We next used Q-DOAS to evaluate the polyamine spermine as an inhibitor of HTT-Ex1(Q44) aggregation (Figure 5C). Small, charged molecules have been shown to affect aggregation by modulating the conformation of amyloidogenic proteins to promote or inhibit amyloid formation^70–74^. In previous work we showed that positively charged polybasic peptides effectively inhibited polyQ aggregation^64^. We thus hypothesized that the small molecule polycation spermine, a naturally occurring polyamine with four amine groups could similarly inhibit aggregation.

We tested the effect of increasing concentrations of spermine on HTT-Ex1(Q44) aggregation using both Q-DOAS and ThT assays (Figure 5C). The ThT assay was performed at twice the mHTT-Ex1 concentration of the Q-DOAS assay, resulting in an increased overall rate of aggregation (Figure 5C). Both assays showed significant, concentration-dependent suppression of aggregation by spermine. This suppression manifested primarily at the primary nucleation stage, appearing as a prolonged lag phase prior to entry into an active aggregation phase (Figures 5C, 5D, Extended Data 3B), suggesting that spermine transiently interacts with mHTT-Ex1 monomers to delay nucleation. Importantly, upon nucleation of seeds and entry into the aggregation phase, aggregates formed at rates comparable to no spermine mHTT-Ex1(Q44) controls. Q-DOAS half-times showed a predominantly logarithmic suppression response to increasing spermine concentrations (Figure 5D). Fitting to an inhibitor dose–response model yielded the equation:

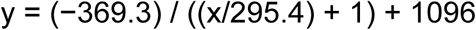

where y is the aggregation half-time in minutes, x is the spermine concentration in µM, and the IC_50_ is 295.4 µM. The fitting was performed using a standard three-parameter inhibitor dose–response model.

Together, these experiments show Q-DOAS is well suited to test protein and small molecule inhibitors of oligomerization and aggregation, and to obtain mechanistic insights into their mode of action. The Q-DOAS readout does not depend on an exogenous dye that can interact non-specifically with small molecules or proteins, although compatibility with potential confounders should be assessed before their inclusion in the assay.

### Detecting HTT seeding activity in cellular and mouse tissue lysates using Q-DOAS

As shown above for αSyn, a potential application of Q-DOAS is the quantification of aggregate-seeding or aggregate-suppressing activity in biological materials. We determined if Q-DOAS could reliably detect seeding-competent material from cellular models expressing normal and pathogenic polyQ-length HTT. HEK293T cells transfected with either HTT-Ex1(Q25)-HA or HTT-Ex1(Q97)-HA to generate HTT aggregates. After cell lysis, aggregates were isolated by sedimentation and detergent washes as in Aviner et al. (2024)^75^*. The aggregates were sonicated to generate seeds and normalized by concentration (Figure 6A). Seeding by aggregates from HTT-Ex1 Q25 and Q97 cells were assessed by Q-DOAS using either mHTT-Ex1(Q44)-BDP (Figure 6B) or ΔN17 mHTT-Ex1(Q44)-BDP (Figure 6C), which is more sensitive to seeding. As expected, Q97-derived aggregates were more seeding-competent than Q25 in both assays (Extended Data 4A,B). QDOAS using mHTT-Ex1(Q44)-BDP displayed increasing fluorescence quenching and shortening the lag phase in a concentration dependent manner (Figure 6A). When ΔN17 was used, the seeding effect and dynamic range of detection was amplified, with bigger effects observed at the lower seed concentrations (Figure 6C). Only the highest concentration of Q97 seeds produces complete quenching over the time course (Figure 6C)*.

**Figure 6.**
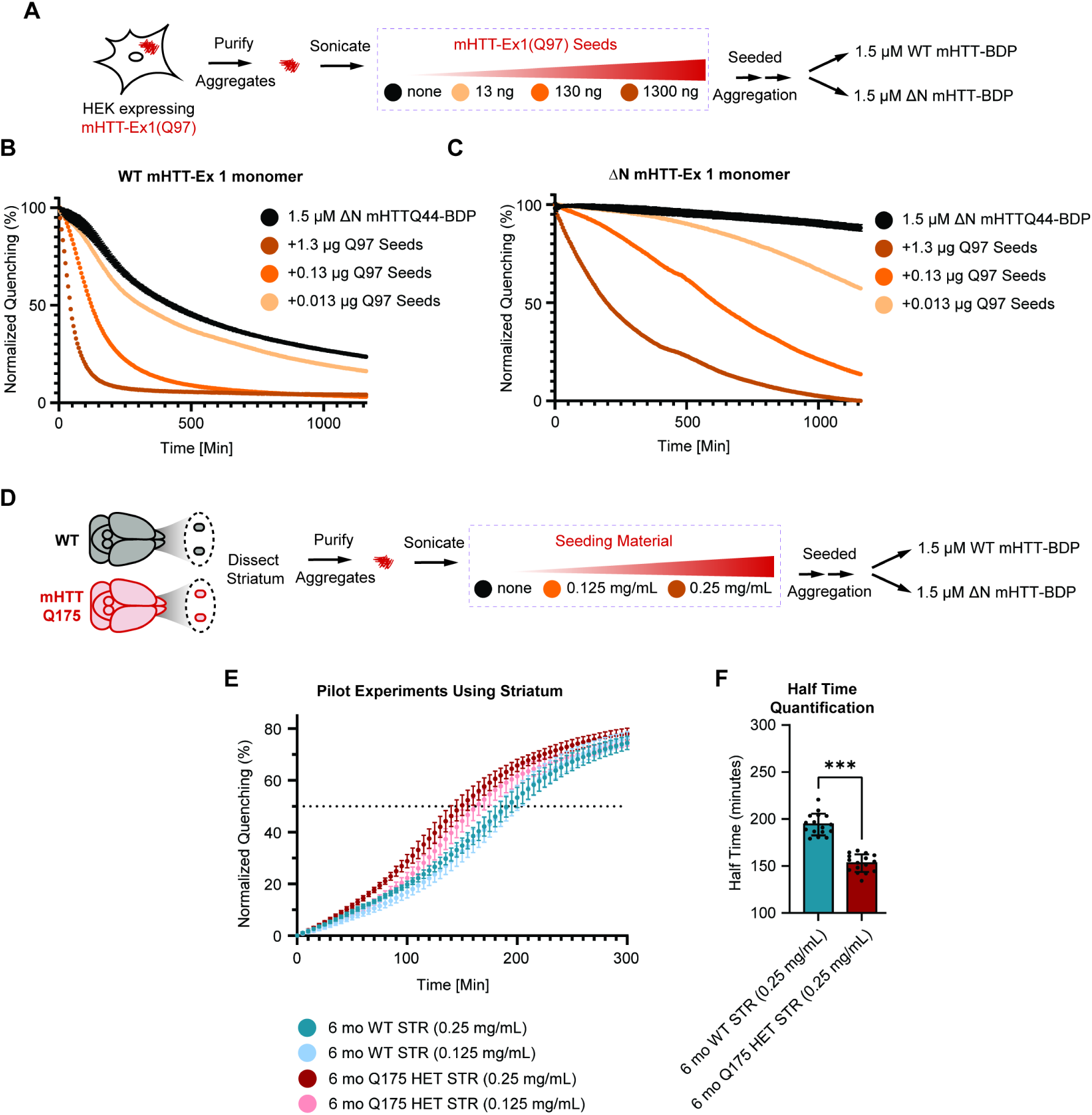
Detecting HTT seeding activity in cellular and mouse tissue lysates using Q-DOAS. (A-C) Q-DOAS experiments using HEK-derived aggregates. (A) Schematic representation of the generation and purification of mHTT-Ex1(Q97) aggregates for addition to Q-DOAS assays using mHTT-Ex1(Q44)-BDP and ΔN17 mHTT-Ex1(Q44)-BDP. (B) Q-DOAS experiments demonstrate a difference between different concentrations of mHTT(Q97)-derived aggregates using mHTT-Ex1(Q44)-BDP. Lysates were made using HEK293T cells and enriched for insoluble seeding compounds. No seed control represents the average of n=2 replicates, whereas seeded curves represent data from n=1 replicate. (C) Q-DOAS experiments demonstrate a difference between different concentrations of mHTT(Q97)-derived aggregates using ΔN17 mHTT-Ex1(Q44)-BDP. No seed control represents the average of n=2 replicates, whereas seeded curves represent data from n=1 replicate. (D-F) Q-DOAS experiments using mouse striatum-derived aggregates. (D) Graphic representation for the preparation of mouse striatum for Q-DOAS analysis. (E) Q-DOAS results using striatum lysate and mHTT-Ex1(Q44)-BDP. All samples show an increase in aggregation over control. Q-DOAS curves shown represent the average of n=16 replicates. (F) Direct comparison of WT/WT and WT/Q175 mouse sample half times show a significant decrease in half time (one-way ANOVA: F = 265.0, P <0.0001; WT vs Q175 P = 0.0006; n=16 per condition).

We next examined if Q-DOAS can detect HTT-seeds in the brain tissue of mouse models of HD^76^. Striata from 6-month-old wild-type and heterozygous HTT-Q175 knock-in mice were dissected, sonicated to release seeding-competent material, and centrifuged to collect the supernatant, which was normalized for Q-DOAS testing (Figure 6D). Addition of HTT Q175-derived lysates increased fluorescence quenching decay relative to WT-derived lysates (Figure 6E), indicating the presence of species promoting mHTT-Ex1 aggregation. Analysis of the half-time of aggregation using Amylofit confirmed a significant increase in aggregation kinetics upon addition of HTT Q175 lysates, relative to WT. The detection of aggregation-promoting species in brain lysates of Q175 mice demonstrates that Q-DOAS can measure the relative potency of aggregation-inducing material in biological samples.

### Design Principles for Adapting Q-DOAS to New Proteins

The successful application of Q-DOAS depends on three key design choices: fluorophore selection, labeling site, and aggregation initiation strategy.

#### Fluorophore Selection

BODIPY-TMR exhibits robust self-quenching when dyes are within 1-2 nm, shows minimal pH sensitivity across physiological ranges, and has favorable photostability for multi-hour kinetic measurements. Alternative self-quenching dyes (e.g., Cy3, Alexa Fluor 555) may be suitable depending on buffer compatibility and spectral requirements. We recommend empirical validation of quenching efficiency by comparing fluorescence of labeled monomers versus pre-formed aggregates.

#### Labeling Site Selection

The labeling site must satisfy two criteria: (1) chemical accessibility for efficient conjugation, and (2) minimal perturbation of aggregation kinetics. Purification and labeling of the recombinant protein of interest are critical steps of the protocol, requiring careful handling to maintain high protein concentration and efficient fluorescent labeling. A labeling site (e.g. a Cysteine) whose chemical modification does not affect aggregation kinetics must be identified. Prior knowledge of amyloid fibril structure and aggregation can guide the choice of a labeling site that will not alter aggregation kinetics^2,5,30,77^. Labeled recombinant protein should be verified by SDS-PAGE and fluorescence measurement to confirm protein purity and labeling efficiency. All buffers should be filtered through a 0.22 µm membrane to minimize contamination. Many buffers contain a reducing agent such as tris(2-carboxyethyl)phosphine (TCEP) or β-mercaptoethanol (βME), and we therefore recommend preparing these fresh. If stored at

−20°C, most buffers may be kept for up to six months. We also recommend subjecting labeled recombinant protein to a 30-minute centrifugation at 30,000 × g immediately before use to pellet any pre-formed aggregates that may arise during thawing. For both mHTT-Ex1 and αSyn, we exploited structural knowledge that terminal regions remain disordered and solvent-exposed in fibrils. We recommend:

- Avoid labeling within structured cores or known nucleation interfaces
- Use substoichiometric labeling (10-50% labeled:unlabeled) to minimize potential artifacts
- Validate that labeled protein aggregates with similar kinetics to unlabeled protein by orthogonal assay (filter trap, ThT if applicable)

#### Aggregation Initiation

For proteins requiring solubility tags (e.g., mHTT-Ex1), TEV cleavage provides synchronous initiation but demands rapid plate loading (<5 min). For proteins that aggregate spontaneously (e.g., αSyn), direct dilution from denaturant or temperature shift may be preferable. For seeding experiments, seeds should be normalized by mass, sonicated to uniform size distribution, and added immediately before measurement to prevent premature assembly.

#### Cross-Validation

We recommend parallel validation using at least one orthogonal method (e.g., filter trap for insoluble aggregates, native PAGE for oligomers) during assay development for each new protein. This anchors the quenching readout to defined biochemical species and establishes confidence in assay interpretation.

#### Q-DOAS Experimental Design

Minimal equipment requirements for the full protocol are met by most biochemistry laboratories: affinity resin, a chromatography instrument equipped with a size exclusion column, gel electrophoresis apparatus, tabletop centrifuges, pH meters, liquid nitrogen, and incubators for bacterial protein expression. Fluorescence measurements require a plate reader and a nano-volume spectrophotometer. Setup requires approximately one to two hours of preparation before mixing reagents, followed by five to ten minutes of active pipetting depending on the scale and complexity of the assay. A 16-channel pipette is recommended for loading bulk reagents across the plate to reduce fatigue, increase throughput, and minimize variability. Certain catalytic reagents — most notably TEV protease — impose a time pressure that demands rapid distribution across wells. The final reaction volume per well in a 384-well plate is approximately 70 µL, which we have found optimal for reducing overflow risk while maintaining a strong fluorescent signal with properly labeled protein. Q-DOAS experiments are typically performed in quadruplicate to reduce the impact of artifacts such as air bubbles introduced during preparation, although as few as two replicates can be used.

## Discussion

We developed Q-DOAS, a single-fluorophore proximity-quenching assay that quantifies protein self-assembly in real time. Using mutant huntingtin exon-1 (mHTT-Ex1) and α-synuclein (αSyn), we demonstrate that Q-DOAS reports oligomerization earlier than Thioflavin T (ThT), yields kinetic data compatible with mechanistic modeling, and quantifies seeding activity in complex biological samples, including patient cerebrospinal fluid and mouse brain tissue.

Plate-based methods for tracking protein aggregation have historically focused on the formation of ThT-reactive amyloid. The ThT assay has served as the gold standard for amyloid detection since its introduction, enabling quantitative study of aggregation-promoting or aggregation-suppressing compounds across a wide range of amyloid-forming proteins^9,30,64,78–81^. However, ThT reactivity depends on the formation of β-sheet-rich assemblies, rendering unstructured or α-helical aggregates invisible to ThT-based readouts^8,41,42^. Researchers wishing to characterize the complete aggregation reaction — including pre-amyloid species — have therefore had to rely on multi-method in vitro approaches or on cell-based assays^82^. Notably, Q-DOAS also avoids a known limitation of ThT: the susceptibility of ThT fluorescence to interference by non-aggregating proteins in complex samples that can bind ThT and small molecules that absorb at ThT excitation or emission wavelengths, or that compete with ThT for binding to fibrils ^10^. Because Q-DOAS does not employ an exogenous dye that must bind to aggregates, this class of artifact is eliminated.

The Q-DOAS assay was developed to enable direct observation of early aggregation events and oligomerization that precede ThT reactivity. Like other FRET-based methods for studying aggregates or protein folding^17,83^, Q-DOAS relies on proximity-dependent fluorescence change to generate an interpretable signal. It distinguishes itself from conventional donor–acceptor FRET approaches by relying exclusively on a single species of self-quenching dye, which eliminates the variability in dye distribution that can arise when two distinct fluorophores are used. BODIPY dyes are also known for their relative insensitivity to changes in pH^84^, supporting a wide range of buffer conditions for studying the biophysical impact of solvent on aggregate formation. Taken together, these properties allow Q-DOAS to track altered aggregation across a broader spectrum of conditions with fewer assay-intrinsic artifacts, improving confidence in the results.

Two proteins with contrasting aggregation behaviors are highlighted here as worked examples: mHTT-Ex1 and αSyn. αSyn exhibits slow aggregation kinetics, whereas mHTT-Ex1 requires an MBP solubility tag that must be cleaved immediately before the assay to prevent premature seeded aggregation. For mHTT-Ex1, fluorescence quenching coincides with oligomer formation detected by native PAGE, occurring hours before SDS-insoluble aggregates appear in filter trap assays. For αSyn, this temporal offset is even more dramatic, with Q-DOAS detecting assembly more than 40 hours before ThT signal rises above baseline. This early detection window provides direct kinetic access to primary nucleation and oligomerization, phases of the aggregation cascade that are largely inaccessible to ThT and are critical for understanding the origin of toxic species. Using this method, we have also been able to explore overall aggregation in mHTT-Ex1 and αSyn in the presence of different buffer conditions and aggregation inhibitors. mHTT-Ex1(Q44)-BDP exhibited significant differences in response to pH and seeding, suggesting that we can observe near monomer-level suppression of HTT aggregation. Notably, prior NMR analyses showed monomeric HTT adopts a less-aggregation prone conformation at low pH^55^, with significant delays in aggregation observable from the beginning of the assay at pH 5.0 (Extended Data 2B). Depending on the protein and the experimental question, the assay can be adapted to address a broad range of questions related to protein aggregation.

The quantitative, time-resolved data from Q-DOAS are directly compatible with kinetic modeling frameworks like AmyloFit, enabling deep mechanistic analysis. We demonstrated this by dissecting the roles of the mHTT-Ex1 flanking domains. Our cross-seeding experiments revealed that aggregation kinetics are dominated by the monomer substrate, not the seed template. The extreme sensitivity of the ΔN17 mutant to seeding, contrasted with the insensitivity of the ΔPRD mutant, kinetically supports a model where the N17 domain promotes primary nucleation—a step bypassed by seeding. This ability to extract quantitative parameters from kinetic profiles provides a powerful tool for dissecting complex aggregation pathways.

Identifying aggregation inhibitors is a key therapeutic goal, but screening is often hampered by assay artifacts. Q-DOAS provides a robust platform for this purpose, as demonstrated with both a protein-based inhibitor (anti-N17 scFv) and a small molecule (spermine). The assay not only quantified inhibition but also provided mechanistic insight, distinguishing the scFv’s role as an oligomerization blocker from spermine’s function as a nucleation inhibitor based on their distinct kinetic signatures. Furthermore, the assay’s sensitivity revealed an unexpected, concentration-dependent inhibitory effect of free GFP on mHTT-Ex1 aggregation, a finding with important implications for the design and interpretation of studies using GFP-fusion proteins. Generally, we see Q-DOAS can quantitatively measure the suppression of overall HTT aggregation and afford researchers an opportunity to measure suppression specifically and at-scale. As the dye can be replaced with any quenchable dye, the method can be fine-tuned to allow for rapid screening of many different small molecules and peptides with high-sensitivity and minimal interaction with the molecule-of-interest.

Q-DOAS can also serve as the basis for quantitative diagnostic methods for amyloid diseases by tracking aggregation competence over time. The Seed Amplification Assay (SAA) represents a technical breakthrough in enabling reliable diagnosis of amyloid diseases from cerebrospinal fluid and other biofluids^49,85^. SAA currently relies on ThT detection of amplified amyloid fibrils, and is therefore insensitive to seeding-competent species that are oligomeric or that exhibit lower amyloid seeding efficiency. Q-DOAS seeks to address this gap by providing quantitative tracking of oligomeric seeding-competent species in biological material, potentially enabling both researchers and clinicians to monitor increases in pre-amyloid aggregation burden over time. We demonstrated its ability to detect seeds across a range of biological complexities, resolving seeding activity in crude lysates from HTT-expressing cells and Q175 mouse striatum. Most notably, the assay detected seeding activity in unamplified CSF from Parkinson’s disease patients, distinguishing them from controls based on aggregation half-time. Future studies could improve the sensitivity of Q-DOAS, for instance by incorporating one or more seed amplification cycles or by including a seed capture step While further optimization and validation will be necessary to demonstrate clinical utility, the ability to derive a quantitative measure of seeding potency from patient samples without amplification suggests this assay could be a promising tool for developing biomarkers to track disease progression or therapeutic response.

### Limitations and Future Directions

Like any biophysical method, Q-DOAS has interpretive boundaries; it reports on proximity, not structure, necessitating orthogonal validation for detailed mechanistic studies. The assay’s performance also depends on careful selection of the labeling site to avoid perturbing aggregation. While we demonstrate proof-of-principle for detecting seeding activity in patient CSF, translation to a clinical diagnostic requires extensive validation in larger cohorts and likely optimization by optimization of the proximity quenching dye of choice. Future work will focus on adapting Q-DOAS to other key amyloidogenic proteins, such as Tau and Aβ, and exploring its use in high-throughput screening campaigns. In conclusion, Q-DOAS provides a versatile, simple and scalable platform to study the earliest events in protein aggregation, addressing a critical methodological gap in amyloid disease research and enabling the real time quantitative measure of aggregation processes.

## Methods

### Expression and Purification of Proteins

#### uTEV3 protein was expressed and purified as described previously^86^

pMAL-MBP-mHTT-Ex1(Q44)-Cys-6xHis plasmids were constructed as described previously^6,87^. Proteins were expressed in Rosetta 2(DE3) pLysS competent cells in 1L TB media supplemented with 1 ng/mL carbenicillin and 0.1 ng/mL chloramphenicol. Cultures were grown to an OD of 0.6 and induced with 1 mL 1 M IPTG (1 mM IPTG final) with 180 RPM shaking overnight at 16°C. 1L cultures were centrifuged at 3,000 ✕ *g* for 30 minutes at 4°C. TB media was decanted and pelleted cells were washed in 25 mL 1X PBS with 1 mM PMSF. Resuspended pellet was transferred to a 50 mL conical tube and centrifuged for another 3,000 ✕ *g* for 30 minutes. 1X PBS was decanted and snap-frozen in LN2 before storage at -80°C until needed for purification. Pellet was resuspended in HTT Lysis Buffer (50 mM HEPES, pH 7.0; 600 mM NaCl; 50 mM imidazole; 1 mM PMSF; 2 mM β-mercaptoethanol; cOmplete™ EDTA-free Protease Inhibitor Cocktail (Roche); 250 units benzonase (Millipore)) and pressure lysed; we have accomplished this by using an Emulsiflex (Avestin). Lysate was cleared for 30 minutes at 20,000 ✕ *g* at 4°C and supernatant was incubated with 10 mL packed Ni Sepharose® High Performance (Cytiva) for one hour with gentle agitation. Sample was washed with 10 CV HTT Wash Buffer (50 mM HEPES, pH 7.0; 600 mM NaCl; 50 mM imidazole; 2 mM β-mercaptoethanol) before eluting protein with HTT Elution Buffer (50 mM HEPES, pH 7.0; 600 mM NaCl; 500 mM imidazole; 2 mM β-mercaptoethanol). Fractions which registered the highest protein concentrations via Bradford assay were collected and concentrated down to 2 mL. Size exclusion was performed with a HiLoad 16/600 Superdex 200 pg (Cytiva) equilibrated with 50 mM HEPES, pH7.0; 500 mM NaCl; 1 mM β-mercaptoethanol; 10% (v/v) glycerol and fractions from the 70 kDa peak were pooled. Pooled fractions were concentrated before LN2 snap-freezing and storage at -80°C. Purity of all proteins was verified using SDS-PAGE and concentrations were determined using the Pierce™ 660nm Protein Assay Reagent (Thermo Scientific). ɑSynuclein A53T A140C plasmids were constructed using Gibson assembly and purified from Rosetta 2(DE3) pLysS competent cells according to preestablished protocols^88^.

### Fluorescent Labeling of Proteins

#### The protocol found here has been published on protocols.io^89^

MBP-mHTT-Ex1(Q44)-Cys-6xHis in 50 mM HEPES, pH 7.0; 500 mM NaCl; 1 mM TCEP; 10% (v/v) glycerol was diluted and heated to RT by allowing the sample to sit out at ambient temperature for 30 minutes. BDP-TMR maleimide (Lumiprobe) was suspended with anhydrous DMSO to a stock concentration of 10000 μM (approximately 192.16 μL DMSO per mg BDP-TMR) and combined with the RT sample at a 10:1 molar excess of dye to protein. Sample was covered with foil and incubated at RT for 2 hours; the reaction was quenched using a 10:1 molar excess of DTT to dye and incubated for another hour at RT. Excess dye was removed using a Zeba™ Spin Desalting Column, 7K MWCO (Thermo Scientific). Size exclusion was performed with a HiLoad 16/600 Superdex 200 pg (Cytiva) equilibrated with 50mM HEPES, pH 7.0; 500 mM NaCl; 1mM β-mercaptoethanol; 10% (v/v) glycerol and fractions from the higher approximately 70 kDa peak were pooled and concentrated. Concentrated fraction was transferred into a siliconized tube and centrifuged at 30,000 ✕ *g* at 4°C for 30 minutes to remove potential aggregates. An aliquot for determining labeling efficiency and concentration was removed before LN2 snap-freezing and storage at -80°C. Optimal fluorescence signal is seen at approximately 55-60% labeling efficiency, with increasing variability and decreasing signal sensitivity as the ratio of dyed to undyed mHTT-ex1 decreases (Figure S1A, S1C-E).

ɑSynuclein A53T A140C was diluted in 1X PBS, pH 7.4; 1 mM TCEP and heated to RT by allowing the sample to sit out at ambient temperature for 30 minutes. BDP-TMR maleimide (Lumiprobe) was suspended with anhydrous DMSO to a stock concentration of 10000 μM (approximately 192.16 μL DMSO per mg BDP-TMR) and combined with the RT sample at a 10:1 molar excess of dye to protein. Sample was covered with foil and incubated at 4°C overnight; the reaction was quenched using a 10:1 molar excess of DTT to dye and transferred into a Pierce™ Slide-A-Lyzer™ Dialysis Cassette, 3.5K MWCO (Thermo Scientific). Excess dye was removed with overnight dialysis in 1X PBS, pH 7.4; 1 mM TCEP at 4°C for 36 hours. Dialyzed fractions were recovered and pooled into a concentration tube. Concentrated fraction was placed into a siliconized tube and centrifuged at 20,000 ✕ *g* at 4°C for 30 minutes to remove potential aggregates. An aliquot for determining labeling efficiency and concentration was removed before LN2 snap-freezing and storage at -80°C. Optimal fluorescence signal is seen at approximately 50% labeling efficiency, with increasing variability and decreasing signal sensitivity as the ratio of dyed to undyed ɑSyn decreases (Figure S1B).

### Q-DOAS Assay (Plate Reader)

The protocol found here has been published on protocols.io^90^.

#### Reaction Buffer A

All reactions are performed in buffer with a final composition of 50 mM HEPES, pH 7.2; 100 mM NaCl; 1 mM TCEP; 3.3% (v/v) glycerol, referred to in the above text as Reaction Buffer A.

#### 16X HTT Q-DOAS Buffer (100 mM Final Salt)

Prepare a 10 mM L-cysteine stock in 10 mL deionized water. Combine 2 mL 1 M HEPES, pH 7.4; 3200 μL 5 M NaCl; 80 μL 500 mM TCEP, pH 7.0; 4 mL 10 mM TCEP; and 1520

μL deionized water (200 mM HEPES, pH 7.4; 1600 mM NaCl; 4 mM TCEP, pH 7.0; 4 mM L-cysteine). Filter the solution through a 0.22 μm filter, aliquot, and store at -20°C for 6 months. Before use, thaw and dilute 16X HTT Q-DOAS Buffer to a 4X stock concentration in deionized water.

#### 16X HTT Q-DOAS Buffer (Salt-Free)

To adjust salt concentration during the Q-DOAS assay, this buffer protocol can serve as an alternative. Prepare a 10 mM L-cysteine stock in 10 mL deionized water. Combine 2 mL 1 M HEPES, pH 7.4; 80 μL 500 mM TCEP, pH 7.0; 4 mL 10 mM TCEP; and 3920 μL deionized water (200 mM HEPES, pH 7.4; 4 mM TCEP, pH 7.0; 4 mM L-cysteine). Filter the solution through a 0.22 μm filter, aliquot, and store at -20°C for 6 months. Before use, thaw and dilute 16X HTT Q-DOAS Buffer to a 4X stock concentration in deionized water.

#### 1X PBS (ɑSynuclein Variant)

Immediately before use, prepare 50 mL 1X PBS solution with 1 mM TCEP. Add 5 mL 10X PBS and 100 μL 500 mM TCEP, pH 7.0 to a graduated cylinder and bring the final volume to 50 mL with deionized water. Filter through a 0.22 μm filter and keep at room temperature until ready to use. Dispose of excess to prevent contamination between assays.

#### mHTT-Ex1-BDP Q-DOAS

1. Place a 384 Well Black Plate with Optically Clear Polymer Bottom (Thermo Scientific, Cat. #: 142761) on a clean benchtop surface. Fill unused wells directly around wells which intend to be used with 70 µL deionized water to reduce evaporation.
  a. **NOTE: Avoid introducing bubbles during preparation.**
2. Thaw and dilute 16X HTT Q-DOAS Buffer (100 mM Final Salt) to a 4X stock concentration. To each well, add 17.5 µL 4X HTT Q-DOAS Buffer (100 mM Final Salt).
  a. If using 16X HTT Q-DOAS Buffer (Salt-Free), supplementary salt will need to be added. To each well, add 17.5 µL 4X desired final salt concentration with 50 mM HEPES, pH 7.4. Should you desire to test additional compounds, combine them with the 4X salt concentration and 50 mM HEPES, pH 7.4 to keep pH and salt concentrations stable.
3. Thaw concentrated mHTT-Ex1-BDP on ice; protect from light by covering the sample on aluminum foil. Dilute in 50 mM HEPES, pH 7.4 to a 4X (6 µM) HTT stock concentration.
4. Centrifuge the 4X (6 µM) HTT stock at 30,000 ✕ *g* at 4°C for 30 minutes to remove potential aggregates. Clean purifications should have very little to no sedimentation visible after the centrifugation. Add 17.5 µL 4X (6 µM) HTT stock to each well.
5. Prepare a 4X stock for each compound of interest at each desired concentration diluted with 50 mM HEPES, pH 7.4. Add 17.5 µL each 4X compound of interest to each respective well.
6. Thaw uTEV3 protease on ice; dilute to a 4 µM 4X concentration. Rapidly add 17.5 µL 4X uTEV3 to each well to begin MBP cleavage. Seal the 384 well plate with optical tape and place in a plate reader set to 30°C with the following settings: Focal Height: 4.7 mm; Flashes per well/cycle: 20; Double Orbital Shaking: 550 rpm; Bottom Optic. Perform gain adjustment using the brightest measured well before beginning the assay; measure fluorescence every 300 seconds (five minutes) until readings have plateaued.
7. Export the raw measurements and import into your preferred data analysis pipeline. To normalize the data in a format amenable to analysis, first transform the data using the formula *y* = 1000000 - *x*, where x is the measured fluorescence value. Data can now be used as it is now for analysis in Amylofit or normalized for analysis via other means.

#### ɑSynuclein Q-DOAS

1. Place a 384 Well Black Plate with Optically Clear Polymer Bottom (Thermo Scientific, Cat. #: 142761) on a clean benchtop surface. Fill unused wells directly around wells which intend to be used with 70 µL deionized water to reduce evaporation.
  a. **NOTE: Avoid introducing bubbles during preparation.**
2. Prepare 1X PBS with 1 mM TCEP as outlined in Reagent Setup (above).
3. Thaw concentrated ɑSyn A53T A140C-BDP on ice to reduce aggregation; protect from light by covering the sample on aluminum foil. Dilute in 1X PBS with 1 mM TCEP to a 2X (60 µM) ɑSyn stock concentration.
4. Centrifuge the 2X (60 µM) ɑSyn stock at 30,000 ✕ *g* at 4°C for 30 minutes to remove potential aggregates. Clean purifications should have very little to no sedimentation visible after the centrifugation. Add 35 µL 2X (60 µM) ɑSyn stock to each well.
5. Prepare a 2X stock for each compound of interest at each desired concentration diluted with 1X PBS with 1 mM TCEP. Add 35 µL each 2X compound of interest to each respective well.
6. Seal the 384 well plate with optical tape and place in a plate reader set to 37°C with the following settings: Focal Height: 4.7 mm; Flashes per well/cycle: 20; Double Orbital Shaking: 400 rpm; Bottom Optic; Idle Double Orbital Shaking: 400 rpm, 30s on, 270s off. Perform gain adjustment using the brightest measured well before beginning the assay; measure fluorescence every 1800 seconds (30 minutes) until readings have plateaued.
7. Export the raw measurements and import into your preferred data analysis pipeline. To normalize the data in a format amenable to analysis, first transform the data using the formula *y* = 1000000 - *x*, where x is the measured fluorescence value. Data can now be used as it is now for analysis in Amylofit or normalized for analysis via other means.

### *In vitro* Aggregation Assay for Generation of Oligomer and Amyloid Seeds

Samples were aggregated according to previous protocols^91^. Mutant Huntingtin aggregation reactions were performed using 1.5 μM of mHTT-Ex1(Q44)-(BDP TMR), 1 μM uTEV3. Aggregation was conducted in Aggregation Buffer (50 mM HEPES, pH 7.2; 50 mM TRIS-HCl, pH 7.2; 0.5 mM EDTA, pH 7.2; 100 mM NaCl; 1 mM TCEP; 3.3% (v/v) glycerol) in 1.5 mL Protein LoBind tubes (Eppendorf) and incubated at 30°C on a rotor. Samples at varied time-points were taken and either snap-frozen in LN2 and stored at -80°C or combined in a 1:1 ratio with a Filter Trap Stop Solution (4% (w/v) SDS, 100 mM DTT), boiled for 5 minutes at 95°C, and stored at -20°C. A separate mutant Huntingtin aggregation was performed using 30 μg of mHTT-Ex1(Q44)-(BDP TMR) and 1 μM uTEV3 in Aggregation Buffer. These samples were allowed to aggregate for 48 hours before centrifugation at 21,000 ✕ *g* at 4°C, resuspension in 1X PBS, pH 7.4, and snap-freezing in LN2.

ɑSyn aggregation reactions were performed using a modified Seed Generation Protocol^47^ with 30 μM ɑSyn A53T A140C-(BDP TMR) in 1X PBS, pH 7.4. In brief, samples were centrifuged at 30,000 ✕ *g* at 4°C for one hour to pellet aggregated material. The supernatant was transferred to a new 1.5 mL Protein LoBind tube (Eppendorf) and placed in a ThermoMixer C (Eppendorf) set to 900 RPM at 37°C and aggregated for 48 hours. Samples were centrifuged at 21,000 ✕ *g* at RT before resuspension in 1X PBS, pH 7.4, and snap-freezing in LN2.

### SDS-PAGE, BN-PAGE and Filter Trap

Aggregated samples processed using the Filter Trap assay were prepared using preestablished protocols^40,92^. SDS-incubated mHTT-Ex1(Q44)-(BDP TMR) samples were filtered through a 0.22 μm cellulose acetate membrane (Sterilitech) backed with gel blotting paper (Whatman) through a constant vacuum and washed with 0.1% (w/v) SDS. Snap-frozen mHTT-ex1(Q44) samples were used for diluted and split for use in either SDS-PAGE^93^ or BN-PAGE^94^ experiments. 10 μL SDS-PAGE samples were combined with 5 μL 4X Laemmli Buffer (300 mM Tris-HCl; 100 mM DTT; 2.4% SDS (v/v); 60% (v/v)) and boiled at 95°C for 5 minutes. 10 μL prepared samples were loaded onto two Novex WedgeWell 4-20% Tris-Glycine gels (Thermo Fisher) with a Precision Plus Protein Dual Color Standard (Bio-Rad) lane and run under 150V current. Gels were either stained using a stain solution (0.1% (w/v) Coomassie R-250; 40% (v/v) ethanol; 10% (v/v) acetic acid) and resolved using a destain (20% (v/v) ethanol; 10% (v/v) acetic acid), or transferred to a nitrocellulose membrane using an iBlot^TM^ 2 Gel Transfer Device (Invitrogen) before Western Blotting.

10 μL native samples were combined with 5 μL 4X Native Sample Buffer (200 mM Bis-Tris, pH 7.0; 60 mM HCl; 10% glycerol (v/v)) and 5 μL Aggregation Buffer. BN-PAGE was performed using a modified version of the original BN-PAGE protocol^94^. In brief, Anode Buffer (50 mM Bis-Tris, pH 7.0), Dark Cathode Buffer (15 mM Bis-Tris, pH 7.0; 300 mM Tricine; 0.2% (wt/vol) Coomassie G250), and Light Cathode Buffer (15 mM Bis-Tris, pH 7.0; 300 mM Tricine; 0.02% (wt/vol) Coomassie G250) were prepared at 4°C. A NativePAGE™ 3-12% Bis-Tris Protein Gel (Thermo Scientific) was removed and the wells were washed with Dark Cathode buffer. Gels were placed in an electrophoresis box surrounded by an ice slurry. The central cavity was filled with Dark Cathode Buffer, enough to cover the wells, and the surrounding cavity was filled with Anode Buffer. 10 μL native samples were loaded into the Dark Cathode Buffer-filled wells alongside a NativeMark^TM^ Unstained Protein Standard (Thermo Scientific), and samples were run under 150V current until the dye front has traveled 1/3 the length of the gel (approximately 30 minutes). Dark Cathode Buffer was removed and replaced with Light Cathode Buffer before running the gel under 250V current until the dye front has traveled to the end of the gel (approximately one hour). One gel was removed and resolved using destain and the second was removed and prepared for a wet transfer.

Wet transfer was performed using a modified version of preestablished protocols^64^. Gels undergoing a wet transfer were incubated in 0.1% (v/v) SDS for 15 minutes. A PVDF membrane was preactivated using 100% methanol for 30 seconds before rinsing in ddH_2_O. The PVDF membrane was then placed in 1X NuPAGE^TM^ Transfer Buffer (Invitrogen). The transfer sandwich was assembled and placed in a gel transfer box surrounded by an ice slurry and run overnight (16 hours) at 10V. Membrane was removed and fixed in 8% (v/v) acetic acid for 15 minutes. Membrane was washed twice for 10 minutes in ddH_2_O, then once in 100% methanol for 10 minutes. Methanol was removed and membrane was washed twice in deionized water for 5 minutes before proceeding to Western Blotting.

Samples in Filter Trap Stop Solution were processed using the Filter Trap assay following preestablished protocols^64,92^. All Western Blotting membranes were blocked using 5% (w/v) BSA in 1X PBS (RPI) and probed using 1:5000 (v/v) mAB5492 (Sigma-Aldrich) for the mHTT-Ex1(Q44) PRD.

### ThioflavinT Assay

The mHTT-Ex1(Q44) ThT assay was performed using a modified version of preestablished protocols^9^. In brief, 3 μM mHTT-Ex1(Q44), 1 μM uTEV3, and respective concentrations of proteins or small molecules were combined in ThT Reaction Buffer (12.5 μM ThioflavinT; 50 mM HEPES, pH 7.2; 100 mM NaCl; 1 mM L-cysteine; 1 mM TCEP). Replicates were placed in a 384 Well Black Plate with Optically Clear Polymer Bottom (Thermo Scientific) and sealed using optical tape. The plate was placed within a plate reader set to 30°C and measured with the following settings: Interval Time: 15 minutes; Excitation: 446 nm; Emission: 490 nm; Focal Height: 4.7 mm; Flashes per well/cycle: 20; Double Orbital Shaking: 550 rpm; Bottom Optic.

ɑSyn A53T A140C-(BDP TMR) was assessed with ThT using alternate settings. In brief, 30 μM ɑSyn A53T A140C-(BDP TMR) and respective concentrations of proteins were combined in ɑSyn ThT Reaction Buffer (12.5 μM ThioflavinT; 50 mM HEPES, pH 7.2; 100 mM NaCl; 1 mM TCEP). Replicates were placed in a 384 Well Black Plate with Optically Clear Polymer Bottom (Thermo Scientific) and sealed using optical tape. The plate was placed within a plate reader set to 37°C and measured with the following settings: Interval Time: 6.65 minutes; Excitation: 446 nm; Emission: 490 nm; Focal Height: 4.7 mm; Flashes per well/cycle: 20; Double Orbital Shaking: 400 rpm; Idle Double Orbital Shaking: 400 rpm; Bottom Optic.

### Mammalian Cell Culturing, Lysis, and Preparation

HEK293T cells (CRL-3216) from ATCC were cultured as previously described^75,95^. In brief, 5✕10^5^ HEK293T cells were grown at 37°C in a 5% CO_2_ incubator in 6-well poly-d-lysine-coated plates in DMEM high-glucose medium (Thermo Fisher). Cells were transfected with pCDNA3 HTT_ex1_(Q25)-HA and pCDNA3 HTT_ex1_(Q97)-HA using Lipofectamine 2000 (Thermo) according to the manufacturer’s instructions. Media was aspirated and HEK293T cells were washed twice in 10 mL 4°C 1X PBS with 1 mM PMSF. Cells were pelleted at 800 ✕ *g* for 10 minutes and resuspended in 1 mL 4°C 1X PBS with 1 mM PMSF before pelleting again at 800 ✕ *g* for 5 minutes. Supernatant was removed and pellets were snap-frozen in LN2 and stored at -80°C until lysis.

Pellets were resuspended in 1 mL Lysis Buffer (50 mM HEPES, pH 7.4; 100 mM NaCl; 0.5% (v/v) NP-40; 1 mM PMSF; 1X Benzonase; 1X Protease Inhibitor Cocktail) and sonicated at 20% amplitude for five 1 s-on/4 s-off cycles on ice. Lysates were centrifuged at 800 ✕ *g* for 2 minutes to remove cell debris; pellet was discarded and preparation proceeded with supernatant. Concentrations were determined using the Pierce™ 660nm Protein Assay Reagent (Thermo Scientific) and supernatants were diluted in Lysis Buffer to equivalent concentrations; volume was noted here for final resuspension. Samples were pelleted again at 21,000 ✕ *g* for 30 minutes at 4°C and the supernatant was removed. Pellet was resuspended in Urea Buffer (8 M urea; 100 mM NaCl; 100 mM HEPES, pH 7.4) and pelleted again at 21,000 ✕ *g* for 30 minutes at 25°C before removing supernatant; this step was repeated one more time. Urea Buffer was replaced with Protein Buffer (500 mM NaCl; 50 mM HEPES, pH 7.4) and pelleted at 21,000 ✕ *g* for 30 minutes at 4°C. Supernatant was removed one last time before resuspension in Protein Buffer to the same volume noted previously and sonicated at 20% amplitude for five 1 s-on/4 s-off cycles on ice. Aliquots were made and snap-frozen in LN2 before storage at -80°C.

### Animals

Mice were raised according to pre-established protocols^96^. Heterozygous Q175 mHTT knock-in mice carrying mHTT with about 190 CAG repeats were obtained from Jackson Laboratory and aged to designated ages. Animals were housed in standard mouse cages under conventional laboratory conditions, with constant temperature and humidity, 12 h/12 h light/dark cycle and food and water ad libitum. All animal studies were carried out in strict accordance with National Institutes of Health guidelines and approved by the UCLA Institutional Animal Care and Use Committees. The mice were anesthetized by pentobarbital. Brains were removed and carefully dissected to obtain cortical and striatal tissues. Dissected tissues were snap frozen in dry ice and stored in -80°C before further processing.

### Animal Tissue Lysis and Preparation

Dissected tissues were processed using a modified tissue extraction protocol^96,97^. Striatal samples were submerged in Apical Storage Buffer (300 mM NaCl; 50 mM HEPES, pH 7.2; 1 mM TCEP) and homogenized using a plastic microtube douncer. Homogenates were then sonicated at 20% amplitude for two 10 s-on/10 s-off cycles on ice and centrifuged at 3,000 ✕ g for 10 minutes to remove undissolved tissue debris. Supernatants were re-sonicated, an aliquot was removed for whole protein concentration, and the remainder was stored at -80°C until use. Concentrations were determined using the Pierce™ 660nm Protein Assay Reagent (Thermo Scientific).

### Electron Microscopy

Negative Staining was performed using a modified version of preestablished protocols^98,99^. 5 μL of 0.6 μM mHTT-Ex1(Q44) and mHTT-Ex1(Q44)-BDP were applied onto glow discharged 300-mesh carbon-coated copper grids (FDF300-CU, Electron Microscopy Sciences) and incubated for 1 minute. Excess buffer was removed by blotting. For staining, 5 μL of a 1% (w/v) uranyl formate was added on grid for 1 minute before blotting to remove excess, and grids were air dried for approximately 5 minutes. Images were acquired using a Morgagni 100 kV Microscope operated at an accelerating voltage of 100 kV.

### Data Analysis, Statistics, and Data Visualization

Fittings were made by transforming the quenched output about the x-axis and using Amylofit to fit the data using provided fitting parameters (Secondary Nucleation Dominated, Unseeded; *m_0_*: 1.5e-6; *k_+_k_n_* initial: 1e7; *n_c_* initial: 2; *k_+_k_2_* initial: 1e16; *n_2_* initial: 3). Statistical analysis on Amylofit output variables and raw data was performed using GraphPad Prism 10.

## Acknowledgements

We acknowledge Kathleen Poston (Stanford University) and the Knight Initiative for Brain Resilience for providing PD patient CSF and contributive discussions. We acknowledge support by National Institutes of Health through awards T32GM007276 (NIGMS, to K.M.K.) and NS092525 (NINDS, to JF). JF and FUH acknowledge support from the Aligning Science Across Parkinson’s (grant ASAP-000282) initiative administered by the Michael J. Fox Foundation. An earlier version of the manuscript was posted to bioRxiv on 2026-06-16 at https://doi.org/10.64898/2026.06.16.732424.

## Contributions

D.G., K.M.K., and J.F. conceived the project. D.G., K.M.K., S.D., and C.S. contributed to protein expression and purification. L.X.L., X.W.Y. and N.W. contributed to mouse brain tissue dissection. D.G., K.M.K., S.D., and R.C. contributed to Q-DOAS experiments and optimization. K.M.K. contributed to in vitro aggregation assays. C.S. and F.-U.H. contributed to alpha synuclein protein design. K.M.K. and J.A. contributed to Negative Stain experiments. K.M.K. and JF wrote the paper with input from the other authors. These authors contributed equally: Dan Gestaut, Korbin M. Kleczko

## Data availability

Data available on Zenodo under DOI: 10.5281/zenodo.20548672

## Code availability

Only commercially and publicly available code has been used. No new code has been generated for this publication; all data cleaning, preprocessing, analysis, and visualization was performed using CLARIOstar MARS v3.32.101.5 (BMG Labtech), ImageStudio v5.2 (LI-COR), GraphPad Prism v11.0.1.90 (Dotmatics), Matlab R2021a (Mathworks), and Amylofit^3^.

## Extended Data Figure Legends

**Extended Data Figure 1.**
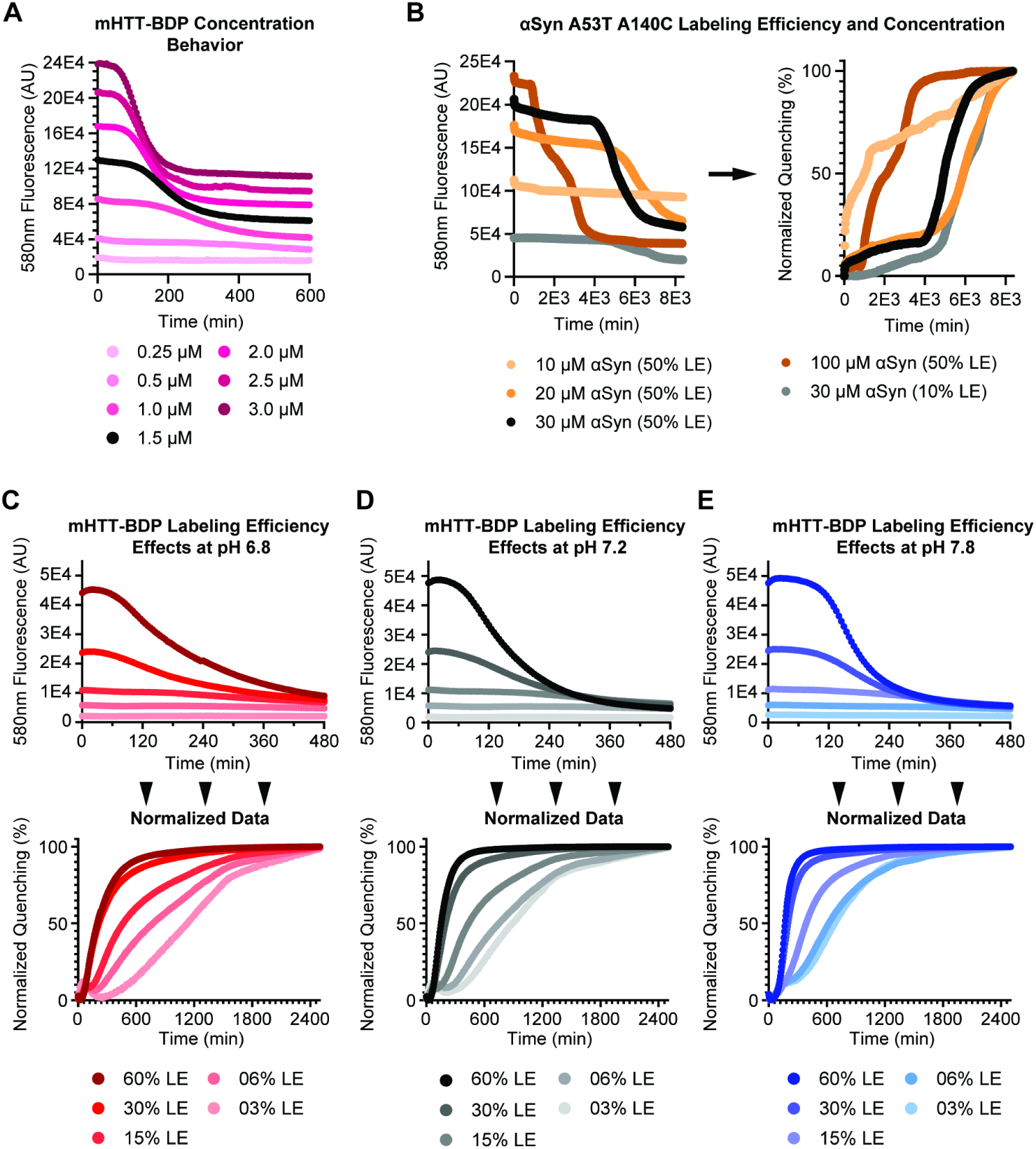
Concentration and labeling efficiency optimization for HTT-Ex1 Q-DOAS experiments. (A) 60% mHTT-Ex1(Q44)-BDP to 40% mHTT-Ex1(Q44) (60% Labeling Efficiency, LE) Q-DOAS was carried out at various concentrations to determine the difference in raw fluorescence signal as concentration of dye and mHTT-Ex1(Q44) in solution are varied. Lower concentrations (below 1.5 µM) exhibit lower sensitivity. Higher concentrations (above 1.5 µM) exhibit higher sensitivity. Curves each represent the data from n=1 replicate. (B) ɑSyn A53T A140C labeling efficiency and concentration experiments exhibit higher variability than mHTT-Ex1(Q44)-BDP. Concentration was increased from 10-100 µM; only 10% and 50% LE were tested; at similar concentration (30 µM), 10% and 50% LE exhibited similar normalized patterns of aggregation, but 10% LE exhibited lower sensitivity. Curves each represent the data from n=4 replicates. (C-E) pH did not significantly affect fluorescence at tested labeling efficiencies, but did show that aggregation was effected regardless of labeling efficiency. (C) Q-DOAS with different BDP-TMR LE at pH 6.8. Curves each represent the data from n=4 replicates. (D) Q-DOAS with different BDP-TMR LE at pH 7.2. Curves each represent the data from n=4 replicates. (E) Q-DOAS with different BDP-TMR LE at pH 7.8. Curves each represent the data from n=4 replicates.

**Extended Data Figure 2.**
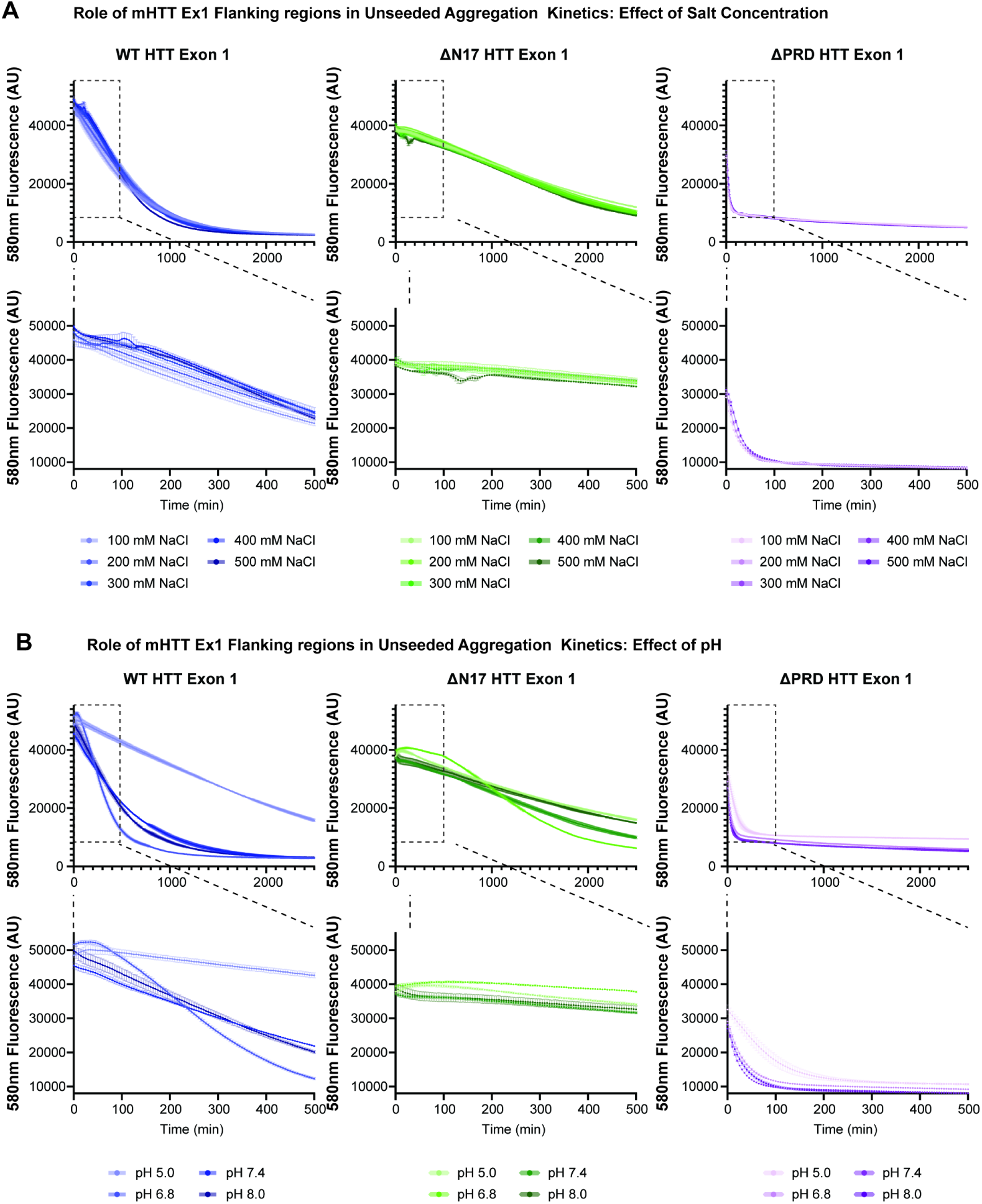
mHTT-Ex1 Q-DOAS Assay sensitivity to pH and salt concentration. (A) Q-DOAS exhibited relatively weak changes in response to increasing salt concentrations from 100-500 mM NaCl across WT, ΔN17, and ΔPRD mHTT-Ex1(Q44)-BDP. (B) Q-DOAS exhibited relatively strong changes in response to increasing pH from 5.0 to 8.0 across WT, ΔN17, and ΔPRD mHTT-Ex1(Q44)-BDP. pH 5.0 exhibited the strongest aggregation suppression, consistent with prior reporting on pH on monomer interactions^55^**. Data represents the average of n=3 replicates.**

**Extended Data Figure 3.**
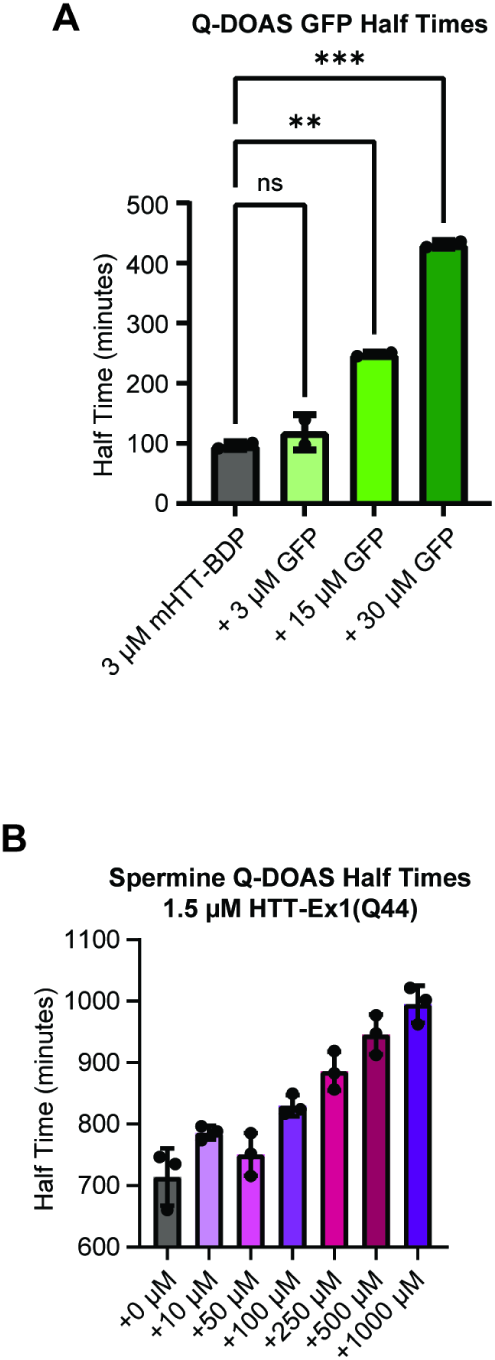
Bar graphs of mHTT-Ex1(Q44)-BDP inhibitor half times. (A) GFP HTT suppression shows a concentration-dependent increase in half-time (one-way ANOVA: F = 197.9, P <0.0001; ** P = 0.0021; *** P = 0.0001; n=2 per condition). (B) Q-DOAS half times from Figure 5 Di were plotted for direct comparison to more clearly see differences at lower concentrations. Data represent the average of n=3 replicates.

**Extended Data Figure 4.**
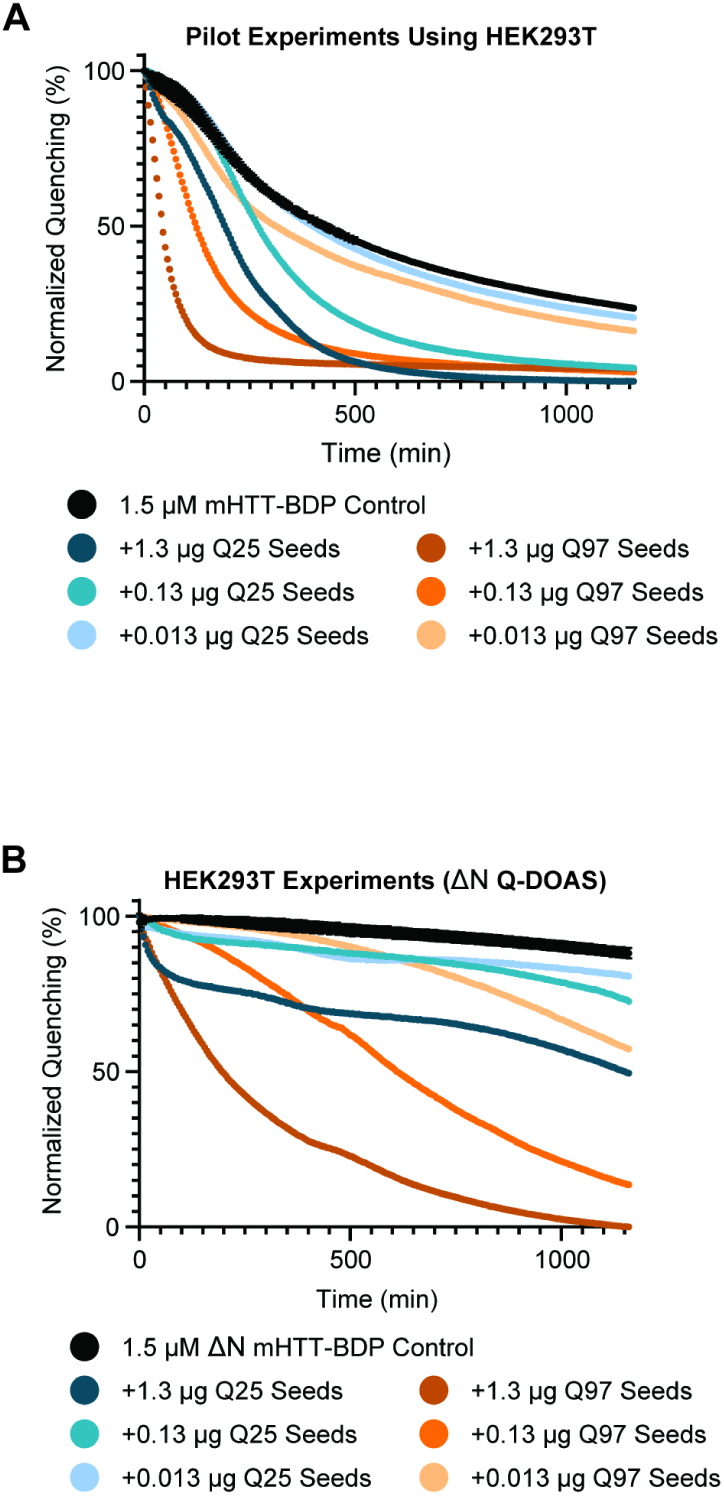
Detecting HTT seeding activity in cellular and mouse tissue lysates using Q-DOAS. (A) Q-DOAS experiments demonstrate a difference between HTT(Q25) and mHTT(Q97)-derived aggregates using mHTT-Ex1(Q44)-BDP. Lysates were made using HEK293T cells and enriched for insoluble seeding compounds. No seed control represents the average of n=2 replicates, whereas seeded curves represent data from n=1 replicate. (B) Q-DOAS experiments demonstrate a difference between HTT(Q25) and mHTT(Q97)-derived aggregates using ΔN mHTT-Ex1(Q44)-BDP. No seed control represents the average of n=2 replicates, whereas seeded curves represent data from n=1 replicate.

